# Altered cortical network in Parkinson’s Disease: the central role of PV interneuron and synaptic remodelling

**DOI:** 10.1101/2025.03.05.641628

**Authors:** Antea Minetti, Elena Montagni, Nicolò Meneghetti, Francesca Macchi, Éléa Coulomb, Alessandra Martello, Alexia Tiberi, Simona Capsoni, Alberto Mazzoni, Anna Letizia Allegra Mascaro, Cristina Spalletti

## Abstract

Parkinson’s disease (PD) is traditionally defined by the progressive degeneration of nigrostriatal dopaminergic neurons; however, accumulating evidence highlights extensive cortical dysfunctions as key contributors to motor and non-motor symptoms. Despite this growing recognition, the precise mechanisms underlying cortical network disruptions and their contribution to PD pathophysiology remain poorly understood, particularly in relation to parvalbumin-positive interneurons (PV-INs) and maladaptive plasticity. Here, we investigate the dysregulation of cortical network homeostasis in PD using a 6-hydroxydopamine (6-OHDA) mouse model, focusing on the progressive disruption of parvalbumin-positive interneuron (PV-IN) connectivity, excitatory/inhibitory balance, and neuroinflammatory responses. Using a multimodal approach integrating longitudinal electrophysiology, wide-field calcium imaging, and histological analyses, we revealed striking alterations in cortical activity and connectivity. Specifically, we observed pathological high-gamma hyperactivity during movement, accompanied by severe disruptions in PV-IN connectivity across motor and somatosensory cortices. Histological analyses further revealed synaptic imbalances and microglial dysregulation, suggesting an extensive cortical response to dopaminergic loss. These findings indicate that PV-IN dysfunction drives cortical maladaptive plasticity, leading to network desynchronization and motor deficits. By reframing PD as a disorder of cortical network homeostasis, this study provides novel mechanistic insights and identifies cortical plasticity as a promising therapeutic target for disease modification.

## INTRODUCTION

Parkinson’s disease (PD) is a chronic and progressive neurodegenerative disorder that primarily affects the motor system, impacting millions of individuals worldwide [1]. The hallmark of PD is the degeneration of dopaminergic neurons in the substantia nigra pars compacta (SNc) and the subsequent depletion of dopamine in the striatum. This dopamine deficiency disrupts the basal ganglia circuitry, leading to the characteristic motor symptoms of PD, including tremors, rigidity, bradykinesia, and postural instability [1–3].

Emerging evidence underscores the central role of dysfunctions within the basal ganglia-thalamo-cortical network in PD pathology. This network is critical for motor control, and its alterations profoundly affect the motor cortex, a region essential for voluntary movement and a promising target for therapeutic interventions [4–6]. A key feature of network dysfunctions is the disruption of cortical and subcortical oscillatory activity. Gamma-band oscillations, in particular, are crucial for motor planning and execution, as they facilitate neuronal synchronization and communication within the motor cortex [7]. Notably, PD patients exhibit a marked reduction in gamma-band oscillations within the basal ganglia-thalamo-cortical network, which correlates with motor dysfunction and symptom severity [8].

The generation and maintenance of gamma-band oscillations depend heavily on GABAA receptor-mediated inhibition, primarily governed by parvalbumin-expressing interneurons (PV-INs) [9]. PV-INs are fast-spiking GABAergic cells that provide strong perisomatic inhibition to pyramidal neurons, regulating cortical network activity, excitatory-inhibitory (E/I) balance, and synchronized oscillatory rhythms [10–12]. Dopamine depletion has been shown to rapidly reduce PV expression in the striatum, suggesting early synaptic dysfunction in PV-INs [13]. Despite growing evidence linking PV-IN dysfunction to basal ganglia pathology, the impact of dopamine depletion on cortical PV-INs and the mechanisms underlying cortical network remodeling in PD remains poorly understood.

Here, we provide a comprehensive functional and anatomical characterization of cortical dysfunctions in a 6-hydroxydopamine (6-OHDA) mouse model of PD. By combining longitudinal electrophysiological recordings, wide-field calcium imaging, and histological analyses, we investigate how dopaminergic degeneration affects PV-IN network connectivity, excitatory/inhibitory balance, and neuroinflammatory responses.

We revealed heightened synchronization in the delta band in striatum over time and increased high-gamma band modulation in motor cortex during voluntary movements.

This progressive breakdown of cortical network integrity is paralleled by disruptions in the cortical PV-IN network, particularly at later stages of pathology progression. These alterations were associated with synaptic remodeling, excitatory/inhibitory imbalance of cortical vesicular markers and heightened microglial phagocytic activity in the motor cortex.

These findings redefine PD as a network disorder and highlight the potential for cortical plasticity modulation as a therapeutic strategy. Understanding these mechanisms may open new avenues for developing interventions aimed at restoring cortical homeostasis and mitigating disease progression.

## METHODS

### Animals

Adult wild-type male C57BL/6J mice (3–4 months old; RRID: IMSR_JAX:000664) were used for electrophysiological and histological analyses (n=8 CNT and n=13 6-OHDA). Adult homozygous PV-Cre knock-in mice (2-3 months old; RRID: IMSR_JAX:008069), were used for wide-field imaging analysis and histological analyses (n=5 CNT and n=4 6-OHDA). Mice were housed up to four animals per cage under a 12-hour/12-hour light/dark cycle with access to food and water *ad libitum*. All experimental procedures respected the ARRIVE guidelines and the European Communities Council Directive #86/609/EEC were approved by the Italian Ministry of Health Autorizzazione n° 544/2023-PR (Risp. a prot. B4BB8.43).

### 6-OHDA Injection

To induce a Parkinsonian model, intra-striatal injections of 6-hydroxydopamine (6-OHDA) were performed. Mice were anesthetized with ketamine (100 mg/kg) and xylazine (10 mg/kg) and placed in a stereotaxic frame. The right striatum was targeted using the following coordinates relative to bregma: anterior-posterior (A/P) +1.0 mm, mediolateral (M/L) −1.8 mm, and dorsoventral (D/V) −3.2 mm. A Hamilton syringe with a fine glass capillary was used to inject 6 mg/mL 6-OHDA dissolved in ascorbic acid solution (0.02% mg/mL) in sterile saline at a rate of 0.2 μL/min for a total volume of 2 μL [14–16]. Control mice received the same volume of physiological saline and ascorbic acid solution.

### Electrophysiological Recordings

#### Surgery

Stainless steel bipolar electrodes were implanted immediately following the 6-OHDA injection. The striatal electrode, targeting the striatum (CPu), was positioned at the same coordinates as the toxin injection, but at a depth of 4 mm. The cortical electrode was inserted 1 mm posterior to the striatal electrode, maintaining the same mediolateral coordinate. This cortical electrode was 1 mm in length, sufficient to reach the superficial layers of the primary motor cortex (M1). An additional hole was drilled at the center of the occipital bone to facilitate the insertion of a surgical screw, which served both as a ground reference and to provide additional stability to the recording implant. The two bipolar electrodes were then soldered to the pins of a connector. The ground pin was connected to the occipital screw, and a first layer of dental cement (Super Bond CeB, Sun Medical Co., Japan) was applied to secure all components to the skull surface. Once the first layer of cement had dried, a second layer (Paladur, Pala, Germany) was applied to enclose the electrical components. To assure a proper head fixation of the animal, crucial for task performance and signal quality during recordings, an L-shaped metal bar (0.6 g) was also attached to the occipital bone using dental cement Super Bond. Following surgery, glucose and paracetamol were administered, and the mice were allowed to recover. After all surgical procedures, animals were carefully stitched and treated with intra-operative analgesia (tramadol 10 mg/kg) and intramuscular injection of cortisone (Bentelan 0.05 ml) upon waking from anesthesia. As a further analgesic, paracetamol (100 mg/kg) was administered for 4 days post-operation in drinking water. The implant enabled simultaneous recordings of neuronal activity in the striatum and motor cortex of the ipsilateral hemisphere relative to the dopaminergic lesion. Electrophysiological recordings were performed during both rest and motor tasks, beginning in the first week post-lesion and continuing until the fourth week.

#### M-Platform for Forelimb Retraction Task

The M-Platform is a robotic system developed in our laboratory for upper limb exercise in mice [17, 18]. It consists of a linear actuator (Micro Cylinder RCL, IAI Germany), a 6-axis load cell (Nano 17, ATI Industrial Automation USA) with a controlled friction system, and a custom-designed handle fixed to the left forelimb positioned on a precision linear sled (IKO BWU 25-75, USA). One end of the handle is screwed onto the load cell for force transfer to the sensor, while the other end serves as a support for the animal’s wrist [19].

#### Electrophysiological recordings at res

After 24 hours of recovery, mice were acclimated to the head restraining system on the M-Platform. Weekly training sessions were conducted for both resting-state and motor task recordings. For resting-state recordings, mice were head-fixed on the M-Platform, with the left forepaw linked to the load cell but without performing the forelimb retraction task. This configuration minimized voluntary movements, allowing the acquisition of neuronal activity exclusively in the ipsilateral hemisphere during rest and excluding any movements sensed by the M-Platform. The chronic recording setup included two bipolar electrodes for simultaneous local field potential (LFP) recordings in the striatum and motor cortex. Signals were amplified using a DigiAmp (Plexon, USA) and referenced to a ground electrode placed at the cerebellum. Data acquisition was performed using the OmniPlex D Neural Data Acquisition System (Plexon Inc., USA). Resting-state signals were recorded for three minutes in an awake but stationary state.

#### Electrophysiological recordings during voluntary motor task

Following the resting-state recordings, mice performed the forelimb retraction task, during which LFP signals from the striatum and motor cortex were recorded. Each session consisted of 15 forelimb retractions, alternating between passive (device-extended by 10 mm) and active (animal-retracted) movements. A sugar water reward was provided for each successful task completion, contingent on surpassing a defined force threshold. Mice typically mastered the task within 2-3 days. [17] Task friction was adjusted based on individual functional deficits. Electrophysiological signals were recorded throughout the task to capture neural activity in the ipsilateral hemisphere during the motor performance. A high-resolution camera (Zyno Full HD, Trust Netherlands, 25 Hz) was positioned parallel to the coronal plane of the animal, capturing video footage synchronized with load cell force signals. Position and velocity signals were extracted from the video and aligned with force signals using a custom algorithm developed in Matlab (MathWorks, USA) (Spalletti et al., 2014). NeuroExplorer software was used to process power spectral density (PSD) data during the task and LFP signals associated with force peaks. Offline analyses, performed with custom algorithms in NeuroExplorer and Matlab, included PSD computation and synchronization of neural and force signals. This setup enabled detailed analysis of cortical and striatal activity during both resting and motor task conditions, providing insights into the neuronal dynamics associated with the lesion-induced motor deficits.

### Electrophysiological Recordings Data Analysis

#### LFP extraction and spectral analysis at rest

From extracellular recordings during the resting state, local field potentials (LFPs) were extracted by applying a low-pass filter at 200 Hz. The LFP signals were then z-scored prior to spectral analysis to normalize the data. Recordings with excessive signal deflection exceeding ±5 SD of the mean were removed from further analyses. The power spectral density (PSD) of the z-scored LFPs was calculated using the Fast Fourier Transform (FFT) implemented via the Welch method (using the pwelch function in MATLAB). For this analysis, the time window of interest was divided into 10-second sub-windows with 50% overlap to improve spectral estimation. Frequency bands of interest were defined as follows: delta (2.5-5.5 Hz), beta (13-30 Hz), and gamma (30-60 Hz), and their respective power was computed as the average PSD within these frequency ranges. For the resting-state analysis we excluded from the LFPs those signals’ portions contaminated by animal movements.

#### Coherence Analysis at rest

The magnitude-squared spectral coherence between striatal and cortical LFPs was estimated using the mscohere function in MATLAB. This function computes the coherence as the ratio between the magnitude-squared cross-power spectral density of the two signals and the product of their individual PSDs.

#### Cross-correlation of Delta Power Fluctuations at rest

To investigate functional connectivity, we computed the cross-correlation between delta power fluctuations of the striatal and cortical LFPs using the xcorr function in MATLAB. Delta power fluctuations were estimated by applying a Hilbert transform to the delta-filtered signal, and then taking the absolute value of the resulting signal to obtain the envelope of the delta activity.

#### Time-frequency analysis of motor task LFPs

To analyze the temporal evolution of spectral patterns in LFPs during motor tasks, we performed time-frequency decomposition using wavelet analysis. LFP scalograms) were generated via continuous wavelet transform (MATLAB *cwt* function) with the analytic Morse wavelet, setting the symmetry parameter to 3 and the time-bandwidth product to 60 for optimal resolution. The entire LFP recording was subjected to wavelet transformation, and the resulting time-frequency scalograms were segmented into windows spanning from −1500 ms to +500 ms relative to movement onset. For normalization, scalograms were baseline-corrected using a reference period between −1500 ms and −750 ms before movement onset. We specifically examined movement-related power modulation in the delta (2.5-5.5 Hz) and high-gamma (100-150 Hz) frequency bands. These were extracted from the scalograms within the [0, 250 ms] window post-movement onset to capture task-related neural activity.

#### M-Platform parameters extraction for motor task

The M-platform parameter extraction was calculated as previously shown in [17]. Briefly, task-related variables were derived from position, speed, and force signals, including the mean force, calculated as the average force from peaks detected during sub-movements, and the area under the force curve (AUC), which represents the total force exerted over time during movement.

#### Statistical analysis

The statistical significance of power, coherence, correlation, and power modulation was evaluated using a two-way ANOVA. The analysis included two factors: recording weeks and animal group (6-OHDA vs CNT). The tests were applied to datasets containing multiple recordings per week across animals, without averaging data at the animal level prior to statistical analysis. To further explore differences between groups, post-hoc pairwise comparisons were conducted between the two animal groups at each recording week using the Wilcoxon rank-sum test, allowing for non-parametric evaluation of group differences.

### Wide-Field Imaging

#### Surgery

PV-Cre mouse line, in which Cre recombinase is selectively expressed under the control of the parvalbumin (PV) promoter, underwent wide-field imaging. For parvalbumin-expressing interneurons (PV-INs) labeling with GCaMP7f, the Cre-dependent viral construct ssAAV-PHP.eB/2-hSyn1-chl-dlox-jGCaMPf(rev)-dlox-WPRE-SV40p(A) (1.3×10E13 vg/ml, volume: 50µL, Viral Vector Facility, ETH, Zurich) was diluted in 100µL of saline solution. A final volume of 150µL was intravenously injected in the retro-orbital sinus of PV-Cre mice under isoflurane anesthesia at P60 to allow wide-brain infection. [20] Two weeks after AAV injection, mice were injected with either 6-OHDA (n=4) to induce a parkinsonian model or vehicle (n=5) for control conditions. Immediately following the injection, the mice underwent an intact-skull preparation to provide direct optical access to the cortex (modified from Montagni et al., 2024 [21]). The surgery was performed under a cocktail of ketamine/xylazine (100/10 mg/kg i.p.). The skin and the periosteum were removed. Bregma was marked for stereotactic reference. A custom-made aluminum head-bar placed behind lambda was glued to the skull using transparent dental cement (Super Bond C&B – Sun Medical). The exposed skull was then covered with a thin layer of the same cement.

#### Wide-field microscopy setup

Imaging was performed on the intact skull of the awake head-fixed animal using a custom-made microscope setup. To excite the GCaMP7f indicator, a 470 nm light beam (LED, M470L3, Thorlabs) filtered by a bandpass filter (469/17.5 nm, Thorlabs) is directed by a dichroic mirror (MD498, Thorlabs) onto the objective lens (TL2X-SAP 2X Super Apochromatic Microscope Objective, 0.1 NA, 56.3 mm WD, Thorlabs). The fluorescence signals was selected by a bandpass filter (525/50 nm filter, Semrock, Rochester, New York, USA) and collected by a CMOS camera (ORCA-Flash4.0 V2 Digital CMOS camera / C11440-22CU, Hamamatsu). Images are acquired with 9 ms exposure time in External Edge mode at 50 Hz for 6000 total frames with a resolution of 512×512 pixels and a field of view (FOV) of 1.15×1.15 cm (depth 16-bit).

#### Habituation and awake imaging

One week after the surgery mice were acclimated to the head-fixation for 5-4 days (10 min a day/mouse) to gradually reduce anxiety and abrupt movements prior to data collection. Head-fixed imaging sessions were performed at 14, 21 and 28 days post lesion (DPL). Each imaging session involves 5 recordings (180 s-long) of spontaneous cortical activity in awake, resting-state mice.

#### Image processing and data analysis

All analyses were performed in MATLAB. Each imaging session was registered using custom-made software, by taking into account the bregma and λ position. An animal-specific field of view template was used to manually adjust the imaging field daily. To dissect the contribution of each cortical area, we registered the cortex to the surface of the Allen Institute Mouse Brain Atlas (www.brain-map.org) projected to our plane of imaging. For each block, image stacks were processed to obtain the estimates of ΔF/F0. ΔF/F0 was computed for each pixel by using the equation ΔF/F0 = (F – F0)/F0, with F representing fluorescence at a given time and F0 the mean fluorescence [21, 22]. The field of view was then downsampled to 128×128 pixels. Global signal regression (GSR) was applied and a total of 22 ROIs were then selected (11 ROI for each hemisphere, 5×5 pixels) representing key cortical regions. Correlation mapping was done for each subject by computing Pearson’s correlation coefficient between the average signals extracted from each ROI, with that of each other ROI. The single-subject correlation maps were transformed using Fisher’s r-to-z transform and then averaged across all animals. Averaged maps were re-transformed to correlation values (r-scores) for figure purpose. For each mouse, r(CNT)-r(6-OHDA) was calculated and averaged across mice in order to visualize matrices of difference between mice injected with vehicle or 6-OHDA.

The abbreviations and extended names for each areas are as follows: MOs-a, anterior region of secondary motor cortex; MOs-p, posterior region of secondary motor cortex; MOp-a, anterior region of primary motor cortex; MOp-p, posterior region of primary motor cortex; SSp-bfd, primary somatosensory area, barrefield; SSp-tr, primary somatosensory area, trunk; SSp-fL, primary somatosensory area, forelimb; SSp-hl, primary somatosensory area, hindlimb; RSP, retrosplenial cortex; VISa, associative visual cortex; VISp, primary visual cortex. Throughout the text and figures, suffixes L and R were added to denote cortical areas of the left or right hemisphere, respectively (e.g., RSP_L_, RSP_R_).

#### Statistical analysis

Network Based Statistic (NBS) Toolbox in MATLAB was used to statistically assess functional network connectivity. [23, 24] We tested for both significantly higher and lower correlations. Differences were considered significant at p<0.05.

### Schallert Cylinder Test

Schallert Cylinder Test was conducted weekly starting from the first week post 6-OHDA injection to assess disease progression [16]. The Cylinder Test is highly effective for assessing unilateral sensorimotor and motor dysfunctions. Animals are placed inside a Plexiglas cylinder (8 cm in diameter, 15 cm in height) adapted for mice according to the protocol described by Schallert and Tillerson. [25] Each animal is recorded for five minutes using a camera (SMXF50BP/EDC, Samsung, Seoul, South Korea) positioned beneath the cylinder. Videos are analyzed frame by frame to count the number of forepaw touches on the cylinder wall. Specifically, the first paw touch on the cylinder walls and the subsequent placement of the forelimbs back on the cylinder base are recorded [18]. To quantify the number of touches made with the limb contralateral to the injection site, the % Contralateral Forelimb was calculated as the ratio of the number of touches made with the forelimb contralateral to the lesion while the animal climbs vertically within the cylinder to the total number of touches. [26]

### Histology

Mice were deeply anesthetized with chloral hydrate and perfused transcardially with 4% paraformaldehyde (PFA, Electron Microscopy Sciences) in 0.1M phosphate buffer. Extracted brains were post-fixed in 4% PFA for 2 hours, followed by 30% sucrose in phosphate buffer at 4°C. Brains were sectioned coronally at 50 μm using a sliding microtome (Leica, Germany) and maintained in PBS for free-floating immunostaining.

For immunostaining, brain slices were incubated in a blocking solution for 1 hour at room temperature (10% donkey serum; 0.3% Triton X-100 in PBS), treated with primary antibodies, prepared at the proper concentration in 1% donkey serum and 0.2 % Triton X-100 in PBS overnight at 4°C. Following 3 washes in PBS, the sections were incubated for 2 hours at room temperature with the specific secondary antibodies. For nuclei visualization Hoechst dye (#B2883; 1:500; Bisbenzimide, Sigma Aldrich, USA). Immunohistochemical analysis was conducted on the substantia nigra pars compacta and striatum using primary anti-tyrosine hydroxylase (1:500 TH, INVITROGEN PA5-85167) and corresponding secondary antibodies for detection of unilateral dopamine degeneration and confirm lesion presence. Cortical plasticity markers such as parvalbumin, PV (1:500, SYSY 1950004); vesicular glutamate transporter type 1, VGLUT1 (1:500, SYSY 135304); vesicular glutamate transporter type 2, VGLUT2 (1:500, SYSY 135403); and vesicular GABA transporter, VGAT (1:500, SYSY 131005) were also investigated in primary motor cortex, particularly vesicular markers in superficial layer II/III. Microglial cells and phagosomes in the cortex were stained respectively with rabbit anti-Iba1 1:500 (Wako, Osaka, Japan, 019–19741) and rat anti-CD68 1:250 (BioRad, #MCA1957).

#### Dopaminergic lesion analysis

TH^+^ fibers signal was acquired using a Zeiss Axio Observer microscope equipped with a Zeiss AxioCam MRm camera (Carl Zeiss MicroImaging GmbH, Germany). For each tissue slice, images were captured with a 10x objective and stitched into a single composite image using Zen Blue Edition software (Carl Zeiss MicroImaging GmbH, Germany). Relative Optical Density (ROD) analysis was performed on 8-bit converted images using ImageJ software (National Institutes of Health, USA) to quantify the mean fluorescence of dopaminergic fibers in the striatum, based on a calibration curve. Quantification of TH-positive neurons in the substantia nigra was conducted using Neurolucida software (MBF Bioscience).

#### Cortical vesicular markers analysis

High-magnification z-stack images (5 stacks, z-step 0.17 µm) of cortical vesicular markers for excitatory (VGLUT1 and VGLUT3) and inhibitory (VGAT) neurons were acquired using a Zeiss Airyscan Confocal Microscope (Carl Zeiss MicroImaging GmbH, Germany), equipped with a 63× oil objective and a 1.3 digital zoom. The images were processed and converted in 8-bit images with ImageJ software to analyze the mean fluorescence of synaptic marker puncta-ring regions surrounding the cell body, subtracting the mean fluorescence of the cell body as background. Similarly, the mean fluorescence of PV puncta surrounding non-PV cell bodies was analyzed to assess the synaptic activity of PV-INs in cortical areas that were identified through wide-field imaging as hypoconnected in the 6-OHDA mouse model. For each hemisphere, a minimum of three cells were analyzed across three fields of view from three brain sections per animal, resulting in a total of n=216 cells per cortical marker. The mean values for the ipsilateral hemisphere were then normalized to the corresponding contralateral hemisphere, which served as the internal control.

#### PV-IN density and distribution

The density and distribution of PV^+^ cell bodies were analyzed across three cortical regions: M1, M2, and S1BF. PV-IN cell bodies signals were acquired using a Zeiss Axio Observer microscope equipped with a Zeiss AxioCam MRm camera (Carl Zeiss MicroImaging GmbH, Germany). For each region, the entire cortical thickness was subdivided into five equal bins, with Bin 1 representing the most superficial layer and Bin 5 the deepest. In M2, due to anatomical constraints, only the three most superficial bins (Bins 1-3) were analyzed. The number of PV-IN cell bodies within each bin was quantified and expressed as the density of cells per square millimeter (cells/mm²). This allowed for a precise assessment of PV-IN distribution across cortical depths within each brain region.

#### PV-IN morphology

PV-IN morphology was analyzed ex vivo in the motor cortex of PV-Cre mice injected with the Cre-dependent AAV-PHP.eB/2-hSyn1-chl-dlox-jGCaMPf(rev)-dlox-WPRE-SV40p(A) virus, following wide-field imaging. The robust GFP signal expressed by the virus enabled the detailed visualization and analysis of small dendrites and all neurons co-localizing with PV^+^ cells. Confocal z-stacks spanning approximately 40 μm were acquired with a z-step of 0.23 μm, resulting in a final voxel size of 0.31 × 0.31 × 0.23 μm. Images were subsequently processed using the Filament Tracer Tool of IMARIS software (Bitplane) for semi-automatic three-dimensional reconstruction and quantitative morphometric analysis. For each hemisphere and analyzed mouse, approximately 10 PV^+^ cortical neurons per animal were reconstructed, allowing for a detailed assessment of dendritic architecture and cellular morphology.

#### Microglia analysis

For microglial density analysis, 1024×1024 pixel images were acquired with a confocal microscope (Stellaris 8 confocal microscope, Leica microsystems) using an 20x objective (HC PL APO CS2, 20x/0.75 DRY), and pinhole was set to 1 AU. Sequential illumination with 488 nm laser lines was used to detect Iba1. The central plane of the sample in the z-axis was selected for image acquisition. Images were processed through Fiji Image software: microglia somata and areas of brain regions (in µm) were counted by combining the Cell Counter and selection of ROI options. Density was calculated as the number of soma divided by the area, and expressed as n of cells/mm^2^.

### Statistical Analysis

Statistical analyses were performed using Prism software (GraphPad, USA). All values reported are the mean ± SEM (indicated in the caption). For all analyses, α = 0.05. To determine the employment of statistical parametric or non-parametric tests, a normality test has been performed (Shapiro-Wilk test or Kolmogorov-Smirnov test). For behavioral test analysis a two-way repeated measures ANOVA followed by Sidak’s multiple comparisons test was used to compare 6-OHDA vs CNT and for longitudinal intragroup studies. For histological analyzes a two-way repeated measures ANOVA followed by Sidak’s multiple comparisons test was used to compare CNT contra, CNT ipsi, 6-OHDA ipsi, 6-OHDA contra.

### PCA Analysis

PCA analysis was performed on python (sklearn.decomposition.PCA) using as a dataset all the collected parameters for each animal (9 animals, 49 features including histology and functional connectivity data). Each feature’s loading in PC1 was assessed by looking at pca.components_ and ranked.

#### Training of the SVM classifier

To identify potential biomarkers for classifying mice as diseased(6-OHDA) or non-diseased (CNT), we implemented a Support Vector Machine (SVM) classification approach using a leave-one-out cross-validation (LOOCV) strategy. The dataset used was the same as for *PCA Analysis*. Prior to classification, the features were standardized using sklearn.preprocessing.StandardScaler to ensure comparability and optimal performance of the SVM. To rigorously evaluate model performance and ensure unbiased estimation of classification accuracy given the low number of rows, we used LOOCV. In this approach, each mouse sample was sequentially left out as the test set while the remaining samples were used to train the SVM model. This process was repeated until every sample had been used once as the test set. The SVM classifier was thus trained using either the entire dataset (All), only histological parameters for puncta markers (Puncta), only histological parameters for lesion quantification - CPu Th^+^ fibers and SNc Th^+^ neurons - (Lesion), only functional connectivity data (FC). To validate the accuracy of the classification model, we generated a scrambled version of the dataset by randomly shuffling the class labels while keeping the feature data intact. The same LOOCV procedure was applied to this scrambled dataset to establish a baseline for comparison (bootstrap).

## RESULTS

### Disrupted Cortical and Striatal Circuit in the 6-OHDA Mice

To induce a slow and stable loss of dopaminergic neurons and model Parkinson’s Disease (PD), we unilaterally injected 6-hydroxydopamine (6-OHDA) into the striatum of adult mice, while control mice (CNT) underwent sham surgery with vehicle injection. For histological evaluation, tyrosine hydroxylase-positive (TH^+^) immunohistochemistry in substantia nigra pars compacta (SNc) was performed (Supplementary Fig. 1a, b). In line with previous evidence [14], a pronounced loss of TH^+^ neurons was observed in SNc of the injured hemisphere, along with a marked reduction in TH^+^ fibers in the caudate-putamen (CPu) (Supplementary Fig. 1c, d). We then evaluated motor deficits by using the Cylinder Test. Our results confirmed a pronounced asymmetry in forelimbs use in 6-OHDA-lesioned mice, characterized by a reduced use of the paw contralateral to the injection without any recovery over time (Supplementary Fig. 1e, f).

In order to evaluate how dopaminergic loss alteres local oscillatory activity and synchronization between the striatum and motor cortex, a chronic implant was placed for simultaneous electrophysiological recordings from both regions following 6-OHDA stereotaxic injection. Local field potentials (LFPs) were recorded weekly from awake, head-restrained mice in resting-state condition (Fig. 1a), when no forelimb movement was detected by the load cell.

**Fig 1.**
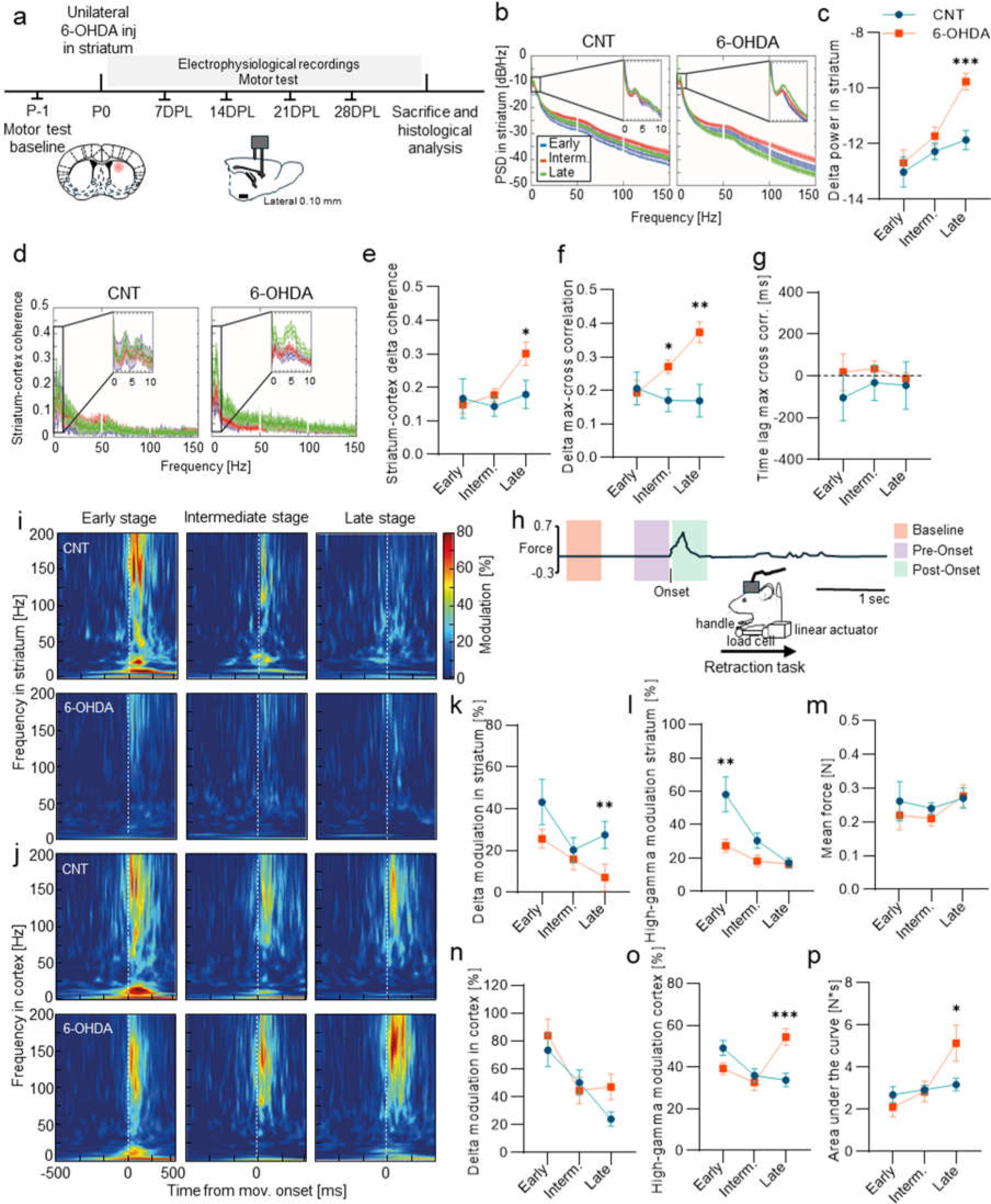
LFPs during resting state revealed increased striatal delta power and striatum-cortical coherence in 6-OHDA-treated mice. **a** The experimental protocol involved unilateral stereotaxic injection of 6-OHDA into the striatum to establish a PD mouse model (in orange n=13), with a control group receiving vehicle injections (in blue n=8). LFPs were simultaneously recorded from the striatum and motor cortex on a weekly basis, during both spontaneous resting states and motor task performance. Recording weeks were grouped into three stages: early (recording week =1); intermediate (2≤recording week≤3); late (recording week =4). **b** Average power spectral density of LFPs in the striatum of control (in blue, CNT n=8) and 6-OHDA mice (in orange, 6-OHDA n=13) across recording stages. Zoomed inset provides focus on the low-frequencies part of the PSD. **c** Spectral power of striatal delta oscillations across recording weeks in controls (in blue, CNT n=8) and 6-OHDA mice (in orange, 6-OHDA n=13). Two-way ANOVA F_StagesXGroups_ = 2.46 p=0.08, F_Stages_ = 10.2 p=0.0001, F_Groups_ 8.51 p=0.004; 6-OHDA vs CNT late stage p=0.0001 Wilcoxon rank-sum test with Holm–Bonferroni correction. **d** Spectral coherence between striatal and cortical LFPs of (in blue, CNT n=8) and 6-OHDA mice (in orange, 6-OHDA n=13) across recording stages. Zoomed inset provides focus on the low-frequencies part of the PSD. **e** Coherence within the delta range in controls (in blue, CNT n=8) and 6-OHDA mice (in orange, 6-OHDA n=13) across recording stages. Two-way ANOVA F_StagesXGroups_ = 2.01 p=0.13, F_Stages_ = 3.82 p=0.02, F_Groups_ = 2.32 p=0.10. 6-OHDA vs CNT late stage p=0.02 Wilcoxon rank-sum test with Holm–Bonferroni correction. **f** Maximal cross-correlation across recording stages of the dynamics of delta power in control (in blue, CNT n=8) and 6-OHDA mice (in orange, 6-OHDA n=13). Two-way ANOVA F_StagesXGroups_ = 3.88 p=0.02, F_Stages_ = 1.9 p=0.15, F_Groups_ = 11.11 p=0.001. 6-OHDA vs CNT intermediate stage p=0.004, 6-OHDA vs CNT late stage p=0.02 Wilcoxon rank-sum test with Holm–Bonferroni correction. **g** Optimal time lag for cross-correlation across recording stages of the dynamics of delta power in control (in blue, CNT n=8) and 6-OHDA mice (in orange, 6-OHDA n=13). Two-way ANOVA F_StagesXGroups_ = 0.14 p=0.87, F_Stages_ = 0.2 p=0.82, F_Groups_ = 1.37 p=0.24. **h** Windows of analysis during the task in relation to movement onset. The black line represents an example of force trace throughout the task; the onset of the movement is highlighted; the orange window identifies the baseline period when no movement is detected; the pink window identifies the pre-onset phase (i.e movement preparation) and the green window the post-onset phase (i.e. movement). **i-j** Average modulation of striatal **i** and cortical **j** LFPs time-frequency scalogram centered around movement onsets (vertical dashed white line) normalized by a baseline ([−1500,−750] ms before movement onsets) across recording stages (indicated in the panels’ titles) in controls (in blue, CNT n=6) and 6-OHDA mice (in orange, 6-OHDA n=7). **k** Modulation of delta power in striatal LFPs around movement onsets (0-250 ms) across recording stages in control (in blue, CNT n=6) and 6-OHDA mice (in orange, 6-OHDA n=7). Two-way ANOVA F_StagesXGroups_ = 0.74 p=0.48, F_Stages_ = 4.29 p=0.01, F_Groups_ = 6.14 p=0.01. 6-OHDA vs CNT late stage p=0.0033 Wilcoxon rank-sum test with Holm–Bonferroni correction. **l** Modulation of high gamma power in striatal LFPs around movement onsets (0-250 ms) across recording stages in control (in blue, CNT n=6) and 6-OHDA mice (in orange, 6-OHDA n=7). Two-way ANOVA F_StagesXGroups_ = 5.02 p=0.007, F_Stages_ = 15.4 p<0.0001, F_Groups_ = 12.31 p=0.0005. 6-OHDA vs CNT early stage p=0.0042 Wilcoxon rank-sum test with Holm–Bonferroni correction. **m** Variation of mean force exerted detected from the robotic M-platform in control (in blue, CNT n=6) and 6-OHDA mice (in orange, 6-OHDA n=7). Repeated Two-way ANOVA Sidak’s multiple comparisons test was used. **n** Cortical delta power modulation in control (in blue, CNT n=6) and 6-OHDA mice (in orange, 6-OHDA n=7): Two-way ANOVA F_StagesXGroups_ = 0.76 p=0.46, F_Stages_ = 8.54 p=0.0002, F_Groups_ = 1.01 p=0.31. **o** Cortical high-gamma power modulation in control (in blue, CNT n=6) and 6-OHDA mice (in orange, 6-OHDA n=7): Two-way ANOVA _StagesXGroups_ = 6.68 p=0.001, F_Stages_ = 3.17 p=0.04, F_Groups_ = 3.99 p=0.04. 6-OHDA vs CNT late stage p=0.001 Wilcoxon rank-sum test with Holm– Bonferroni correction. **p** Variation of area under the force curve detected from the robotic M-platform in control (in blue, CNT n=6) and 6-OHDA mice (in orange, 6-OHDA n=7). Repeated Two-way ANOVA with Sidak’s multiple comparisons test F_StagesXGroups_ (2, 33) = 4.88 p=0.014, F_Stages_ (2, 33) = 6.83 p=0.0033, 6-OHDA vs CNT late stage p=0.037. Mean ± SEM. *p≤0.05; **p<0.01; ***p<0.001; ****p<0.0001. See also Supplementary Figures 1, 2 and Supplementary Table 1.

The LFP of both groups displayed prominent delta oscillations, with a characteristic peak around 4 Hz, in both the striatum (Fig. 1b) and motor cortex (Supplementary Fig. 2a). In the striatum, the power of delta oscillations was modulated by both the recording stage and the experimental group. Over time, 6-OHDA mice demonstrated a clear trend of increasing delta power (Fig. 1c). By the final recording stage, delta power in 6-OHDA mice was significantly higher compared to controls. This progressive increase suggests a temporal evolution of striatal oscillatory activity in 6-OHDA mice. In the motor cortex, delta oscillations in 6-OHDA mice followed a similar upward trend across recording stages (Supplementary Fig. 2a). However, unlike in the striatum, this increase did not reach statistical significance when compared to the corresponding oscillations in controls (Supplementary Fig. 2b).

Across both the striatum and motor cortex, power in other frequency bands, including beta and gamma, did not exhibit clear temporal trends or significant differences between the two groups (Supplementary Fig. 2c-f).

Next, we examined the relationship between the spectral features of LFPs in the striatum and motor cortex across the two experimental groups (Fig. 1d). To evaluate spectral connectivity between these regions, we computed spectral coherence (see Materials and Methods). Both groups exhibited significant delta-band coherence between the striatum and motor cortex during the resting state (Fig. 1e), with coherence being significantly influenced by the recording stage. In the 6-OHDA group, delta coherence progressively and consistently increased over recording stages (Fig. 1e), culminating in a statistically significant difference from controls during the later recording stages.

To further explore this enhanced coupling, we computed the cross-correlation between the temporal dynamics of striatal and cortical delta power (Fig. 1f). Consistent with the coherence findings, cross-correlation peaks demonstrated significant modulation across experimental groups and a significant interaction effect between group and recording week. In 6-OHDA-treated mice, cross-correlation of delta power between the striatum and motor cortex progressively increased over time, becoming significantly elevated compared to control mice as early as the intermediate stage post-toxin injection. These findings indicate that the neurodegenerative processes induced by 6-OHDA injection enhance the synchronization of delta oscillations between the striatum and motor cortex.

Finally, we investigated the time lag at which the maximum cross-correlation occurred between striatal and cortical delta power. No significant differences in the optimal time lag were observed between groups or across recording weeks (Fig. 1g). This suggests the absence of a direct causal relationship in the temporal evolution of delta power between the striatum and motor cortex.

### Voluntary movement induced increased high-gamma spectral modulation in 6-OHDA mice motor cortex

Next, to identify specific alterations in motor function-related activity in the striatum and motor cortex, we investigated the spectral modulation induced by voluntary movement in both control (n=6) and 6-OHDA (n=7) mice. To this end, LFPs were simultaneously recorded from the cortex and striatum as the mice performed a voluntary forelimb retraction task using the M-Platform, a custom-designed apparatus for functional evaluation and neurorehabilitation of forelimb movement in mice [17, 19, 27].

The LFP recordings were segmented into time windows spanning from [−1500 ms to +500 ms] relative to movement onset, which was identified using a custom detection algorithm (see Materials and Methods). Time-frequency scalograms were computed for each segment and normalized to a baseline period, defined as the average scalogram from [−1500 ms to −750 ms], with no force peak detected (Fig. 1h).

In control mice, voluntary movement elicited robust spectral modulation in the striatum, spanning a broad frequency range during the early recording stage (Fig. 1i, top-left). However, this modulation diminished progressively over time, and by the late recording weeks, little to no movement-related spectral activity was observed (Fig. 1i, top-right). In contrast, 6-OHDA mice exhibited minimal movement-induced spectral modulation in the striatum already from the early recording stage, with the only exceptions of delta and high-gamma (Fig. 1i, bottom row). Accordingly, striatal delta power modulation exhibited a decreasing trend in both control and 6-OHDA mice (Fig. 1k), though the reduction was more pronounced in 6-OHDA mice. By the late recording stage, 6-OHDA mice showed a complete loss of delta modulation, which led to a statistically significant difference between the two groups during this phase. Similarly, striatal high-gamma modulation declined progressively in control mice across recording weeks (Fig. 1l). Interestingly, this pattern was not observed in 6-OHDA mice, where high-gamma modulation in the early stage of recording was comparable to the levels seen in control mice at the end of the recording period (Fig. 1l).

A similar progressive reduction in spectral modulation was observed in cortical LFPs of control mice (Fig. 1j, top row). During the initial weeks of recording, voluntary movements elicited increases in power across multiple frequency bands, particularly in the delta, beta, and high-gamma ranges (Fig. 1j, top-left). However, this movement-induced spectral modulation diminished steadily over the course of the recording weeks. In 6-OHDA mice, a similar pattern of movement-induced spectral modulation was observed during the early recording stage (Fig. 1j, bottom-left), with increases in delta, beta, and high-gamma power. Over time, however, spectral modulation in these mice decreased in a manner that paralleled the trend seen in controls, except in the high-gamma range, where an opposing trajectory emerged. Specifically, delta power modulation in the cortex followed a similar declining trajectory in both control and 6-OHDA mice, with a gradual reduction over recording weeks (Fig. 1n). In contrast, high-gamma power modulation displayed divergent trends between the two groups (Fig. 1o). High-gamma progressively decreased in control mice over the recording weeks, while it increased steadily in 6-OHDA mice. By the late stages of recording, this divergence resulted in a statistically significant difference in high-gamma modulation between the two groups (Fig. 1o).

Although significant changes in cortical high-gamma power modulation were observed, these changes did not translate into alterations in mean force output across the recording stages (Fig. 1m). The mean force remained stable in both control and 6-OHDA-treated mice, with no significant differences between the groups. However, the area under the force curve (AUC) increased significantly in 6-OHDA-treated mice during the late stage (Fig. 1p). This increase likely indicates a reduction in the precision of movement, resulting in a more dispersed force profile over time. While the mean force output remains unchanged, its distribution appears less focused, suggesting an alteration in movement control in 6-OHDA-treated mice.

Finally, to further investigate the potential implications of cortical high-gamma modulation in 6-OHDA-treated mice, we assessed its correlation with behavioral and cellular markers (Supplementary Fig. 2g). A significant negative correlation was observed between cortical high-gamma power and performance in the behavioral test at 28 days post-lesion (Supplementary Fig. 2h), indicating that increased cortical high-gamma activity might be associated with motor performance deficits. Additionally, cortical high-gamma power was inversely correlated with the number of TH^+^ cells in SNc (Supplementary Fig. 2i), as well as the intensity of TH^+^ fibres in striatum (Supplementary Fig. 2j), indicating a link between increased cortical high-gamma activity and dopaminergic degeneration. These findings suggest that the pathological increase in cortical high-gamma activity may be the effect of circuit rearrangements in response to progressive nigrostriatal degeneration. Together, these correlations provide a link between cortical spectral changes, cellular degeneration, and motor impairments, highlighting the complex interplay between neurophysiological, behavioral, and cellular adaptations in the 6-OHDA Parkinsonian model.

### Dopaminergic neuron loss impact PV-INs functional cortical network

Electrophysiological results suggest that dopaminergic degeneration may affect broader cortical network connectivity. Specifically, increased high-gamma modulation in the motor cortex of 6-OHDA mice pointed to a compensatory or maladaptive change in neuronal dynamics mediated by parvalbumin-positive interneurons (PV-INs) [10, 28, 29]. Therefore, we used wide-field (WF) calcium imaging to longitudinally investigate how functional cortical connectivity of PV-INs changed from 14 to 28 days post lesion in 6-OHDA mice.

PV-Cre mice were administered retro-orbital injection with a PHP.eB serotype AAV expressing GCaMP7f under the parvalbumin (PV) promoter in a Cre-dependent manner. After two weeks, unilateral stereotaxic injection of 6-OHDA into the striatum was performed to generate the PD mouse model (6-OHDA), with a control group receiving a vehicle injection (CNT). Then, weekly WF imaging and Schallert Cylinder test were conducted starting from 14 days post lesion (DPL) for three weeks (Fig. 2a). The efficiency and specificity of viral infection were validated by post-mortem immunohistological inspection, showing that 90% of PV^+^ neurons were successfully transfected with the GCaMP, and 78% of GCaMP^+^ neurons were effectively identified as PV^+^ neurons (Supplementary Fig. 3a, b). These findings confirm that PHP.eB transfection achieved both high efficacy and target specificity. Moreover, motor deficits in the use of the contralateral forelimb (Supplementary Fig. 3c), dopaminergic neuron loss in both SNc (Supplementary Fig. 3d) and CPu (Supplementary Fig. 3e) were validated, exhibiting a similar trend to that observed in the electrophysiological group (Supplementary Fig. 3d-f).

**Fig. 2.**
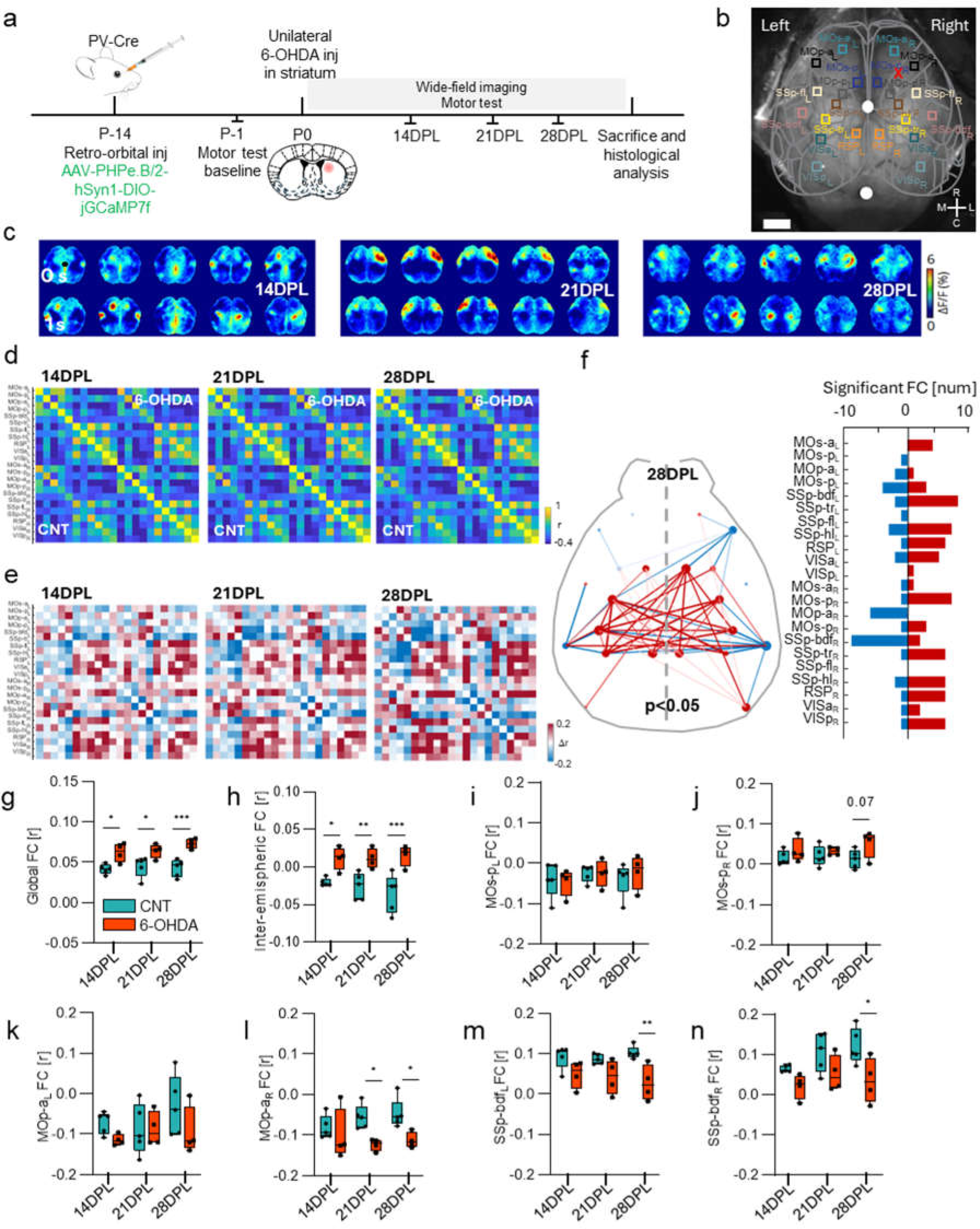
Altered network of PV-INs rs-FC in 6-OHDA mouse model. **a** Experimental timeline for PV-INs longitudinal WF calcium imaging. **b** Representative image of the WF field-of-view. Colored squares represent the cortical parcellation ROI map based on the Allen Brain Atlas. White dots indicate Bregma (top) and Lambda (bottom). Scale bar = 1 mm. **c** Representative sequences of cortical activity in resting state condition (RS) at 14 days post-lesion (14 DPL), 21 and 28 DPL in control (CNT, top) and 6-OHDA treated group (bottom). **d** Dynamics of the averaged functional connectivity PV network (Pearson’s correlation) over the days (14 21 and 28 PDL) for the control group (CNT n=4, bottom right) and 6-OHDA-treated group (6-OHDA n=5, top left). **e** Matrix of difference displaying average FC differences between CNT and 6-OHDA over the days, calculated by subtracting the averaged FC of the CNT group from that of the 6-OHDA group. Red squares represent regions of 6-OHDA hyperconnectivity, while blue squares denote 6-OHDA hypoconnectivity. **f** Network diagram of statistically significant FC alterations at 28 DPL (left). Red and blue lines represent significant hyper- and hypo-correlations of the 6-OHDA group compared with the CNT group, respectively. The bar plot (right) shows the number of significant FC (hypo left, hyper right) related to each cortical area (p<0.05, NBS. n=4 CNT and 5 6-OHDA). **g** Box plot showing the mean global signal PV-FC in the CNT group (light blue) comprate to the 6-OHDA group (orange) at 14,21 and 28 PDL. Repeated Two-way ANOVA with Sidak’s multiple comparisons test (F_Groups_ (1, 7) = 19.12 p=0.003; 6-OHDA vs CNT 14DPL p=0.014; 21DPL p=0.024; 28DPL p=0.0004). **h** Box plot showing the mean inter-hemispheric PV-FC in the CNT group (light blue) comprate to the 6-OHDA group (orange) at 14,21 and 28 PDL. Repeated Two-way ANOVA with 6-OHDA vs CNT (F_Groups_(1, 7) = 19.49 p=0.0031; 6-OHDA vs CNT 14DPL p=0.0250; 21DPL p=0.0094; 28DPL p=0.0005). **i-j** Box plots show comparison of the PV-FC for the posterior area of the secondary motor cortex between CNT and 6-OHDA group (light blue and orange, respectively) at 14, 21 and 28 DPL in the left hemisphere (I, Repeated Two-way ANOVA) and right hemisphere (J, Repeated Two-way ANOVA). **k-l** Box plots show comparison of the PV-FC for the anterior area of the primary motor cortex between CNT and 6-OHDA group (light blue and orange, respectively) at 14, 21 and 28 DPL in the left hemisphere (K,) and right hemisphere (L, Repeated Two-way ANOVA with Sidak’s multiple comparisons test F_Groups_(1, 7) = 13.66 p=0.007; 6-OHDA vs CNT 21DPL p=0.028; 28DPL p=0.033). **m-n** Box plots show comparison of the PV-FC for the barrelfield cortex between CNT and 6-OHDA group (light blue and orange, respectively) at 14, 21 and 28 DPL in the left hemisphere (M, Repeated Two-way ANOVA with Sidak’s multiple comparisons test F_Groups_(1, 7) = 8.81 p=0.021; 6-OHDA vs CNT 21DPL p=0.028; 28DPL p=0.0040) and right hemisphere (N, Repeated Two-way with Sidak’s multiple comparisons test F_Time_ (2, 14) = 4.71 p=0.027, F_Groups_(1, 7) = 7.15 p=0.032; 6-OHDA vs CNT 28DPL p=0.049). In blue, CNT n=5 and in orange, 6-OHDA n=4. Mean ± SEM. *p≤0.05; **p<0.01; ***p<0.001. See also Supplementary Figure 3 and Supplementary Table 2.

In vivo imaging sessions were performed in awake, head-fixed mice at 14, 21 and 28 DPL for both CNT (n=6) and 6-OHDA (n=5) groups (Fig. 2c). Hemoglobin signal was removed from the calcium data, and global signal regression was applied to eliminate global signal contributions. Resting-state functional connectivity (rs-FC) was assessed by defining a set of 22 regions of interest (ROIs) covering both the hemispheres (Fig. 2b). Correlation matrices were computed for each group using pairwise Pearson correlation (Fig. 2d). Differences in rs-FC between CNT and 6-OHDA groups were then quantified by comparing the average correlation matrices of injured mice to those of healthy controls at each time point (Fig. 2e). Results revealed abnormalities in the PV-INs cortical functional network, showing both hypo- and hyper-connectivity patterns in 6-OHDA mice, which appeared to gradually intensify over time (Fig. 2e). Using the network-based statistic (NBS) we therefore tested for significant changes in FC. Relative to the control group, 6-OHDA mice exhibited pronounced hyper-connectivity of the somatosensory areas. Decreased connectivity was instead observed bilaterally in the barrelfield cortex and predominantly in the primary motor cortex of the ipsilateral hemisphere to the lesion (Fig. 2f). Our results showed the emergence of a significantly altered network at 28 DPL, characterized by concurrent patterns of hypo- and hyper-connectivity across brain regions.

To identify the features contributing most to the network alterations, we studied FC dynamics globally and then we further categorized FC into inter-hemispheric or intra-hemispheric connections (Supplementary Fig. 3f). The 6-OHDA group consistently exhibited higher global FC values compared to the control group across all time points (Fig. 2g), suggesting a possible hypersynchronization of the circuitry. These differences increased over time, despite the control group remaining stable throughout, by reflecting an early and acute response to the dopaminergic neuron loss. Interestingly, intra-hemispheric connectivity both contralateral and ipsilateral to the lesion did not show any significant variation over time by highlighting that functional connections within a single hemisphere were more stable and less affected by the lesion (Supplementary Fig. 3g, h). In contrast, transcallosal communications were significantly disrupted among cortical regions in 6-OHDA mice by contributing substantially to the network alterations over all time points investigated (Fig. 2h).

To further evaluate FC alterations potentially related to motor or sensory deficits, we then focused on three main regions of interest: the secondary and primary motor cortex, critical for movement regulation, and the barrelfield cortex, a key area for sensory processing in mice. The secondary motor cortex (MOs-p) showed stable FC over time in both contralateral- and ipsilateral hemisphere (Fig. 2i, j) by suggesting that this cortical region remains relatively resilient to dopaminergic neuron loss. A similar trend was observed also in the primary motor cortex of the contralateral hemisphere (Fig. 2k). In contrast, FC was significantly reduced as early as 21 DPL in the primary motor cortex of the ipsilateral hemisphere (Fig. 2l) indicating an early onset of alterations on motor regions. Barrel field cortex instead exhibited a trend toward hypoconnectivity in the 6-OHDA group with more pronounced effects in the contralateral hemisphere compared to the ipsilateral one (Fig. 2m, n).

Taken together, these results suggest that while the pathological phenotype remains consistent over time, with an immediate and stable impairment, FC shows a progressive deterioration with emergence of an altered network at 28 DPL. Notably, dopaminergic loss affects primarily inter-hemispheric connections by suggesting a stronger impact on the coordination between the two hemispheres.

### Synaptic Puncta Output Compensates for PV-INs Deficiency in the Hypoconnected Areas

In the 6-OHDA mouse model, motor and sensory cortical areas exhibit disrupted functional connectivity of PV-IN networks. PV-INs play a critical role in synchronizing activity across cortical regions and generating gamma oscillations, which are essential for proper cortical processing. This dysfunction may underlie motor and cognitive deficits associated with Parkinson’s disease [30].

Given their role in cortical synchrony and interhemispheric coordination, we investigated potential alterations in PV-IN density, distribution, and synaptic connectivity. By analyzing PV+ neurons and the surrounding puncta rings on non-PV neuronal somata in superficial layers II/III (Fig. 3a), we identified changes in the inhibitory network associated with dopaminergic loss.

**Fig. 3.**
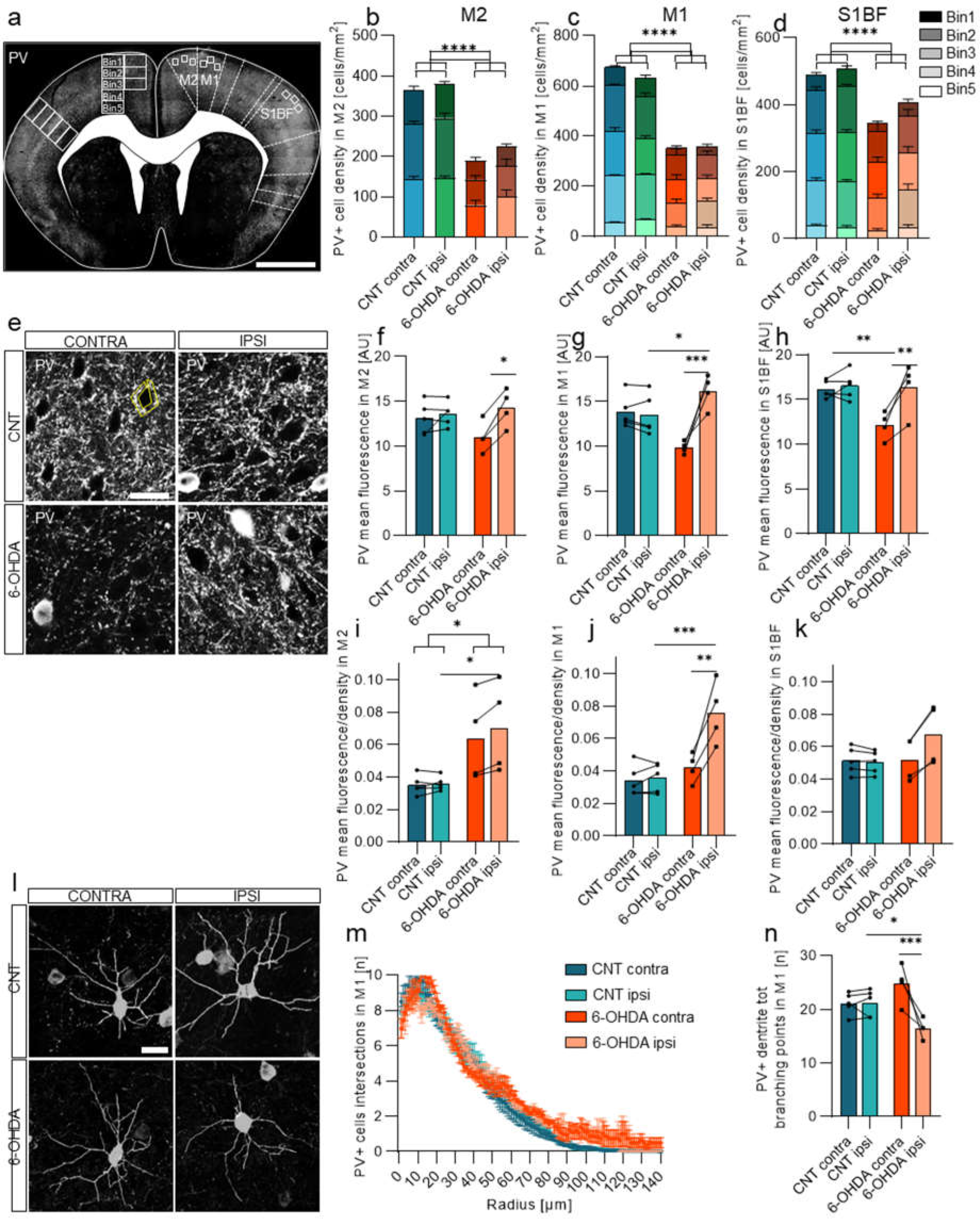
Parvalbumin-positive interneurons (PV-INs) cortical layers’ distribution, density and PV ring point in the cortex. **a** Cortical PV-IN density and layer distribution. Representative coronal section representing the regions of interest. Scale bar: 1 mm. The cortical distribution of PV-INs across five homogenous bins spanning the entire cortical thickness in **b** secondary motor cortex (M2, Two-way ANOVA F_Groups_ (3, 42) = 30.14 p<0.0001, F_Bins_ (2, 42) = 28.40 p<0.0001), **c** primary motor cortex (M1, Two-way ANOVA F_GroupsXBins_ (12, 70) = 2.24 p=0.019, F_Groups_ (3, 70) = 48.09 p<0.0001, F_Bins_ (4, 70) = 69.29 p<0.0001) and **d** primary somatosensory barrel cortex (S1BF, Two-way ANOVA F_Groups_ (3, 70) = 8.91 p<0.0001, F_Bins_ (4, 70) = 66.86 p<0.0001). In particular, in M2 only the 3 more superficial bins have been analyzed due to anatomical regions. Specific statistical values are reported in supp table 5. **e** High magnification of the cortex immunostained with anti-PV for CNT and 6-OHDA groups. Interhemispheric analysis of anti-PV fluorescence mean values of puncta-rings, calculated in puncta-rings around cell bodies of non-PV positive neurons in control and 6-OHDA group. Data from three fields of view per three brain sections n=72 cells for each dataset and CNT animals n=6 vs 6-OHDA animals n=5 were analysed. **f** In M2 interhemispheric analysis of anti-PV fluorescence mean values of puncta-rings in control and 6-OHDA group. Two-way ANOVA with Sidak’s multiple comparisons test F_Hemisphere_ (1, 14) = 5.28 p=0.038; contra vs ipsi 6-OHDA p=0.036. **g** In M1. Two-way ANOVA with Sidak’s multiple comparisons test F_GroupsXHemisphere_ (1, 14) = 14.80 p=0.0018, F_Hemisphere_ (1, 14) = 11.92 p=0.0039; contra vs ipsi 6-OHDA p=0.0005); 6-OHDA vs CNT contra p=0.011. **h** In S1BF. Two-way ANOVA with Sidak’s multiple comparisons test F_GroupsXHemisphere_ (1, 14) = 4.92 p=0.044, F_Groups_ (1, 14) = 6.18 p=0.026, F_Hemisphere_ (1, 14) = 7.45 p=0.016; contra vs ipsi 6-OHDA p=0.010. 6-OHDA vs CNT contra p=0.010. **i-k** Normalized PV puncta density (PV puncta per PV-IN) in superficial layers of the ipsilateral and contralateral hemispheres across cortical regions in the controls (in blue) and 6-OHDA (in orange) group. **i** In M2. Two-way ANOVA with Sidak’s multiple comparisons test F_Groups_ (1, 14) = 12.79 p=0.0030; 6-OHDA vs CNT contra p=0.030. **(j)** In M1. Two-way ANOVA with Sidak’s multiple comparisons test F_GroupsXHemisphere_ (1, 14) = 7.93 p=0.014, F_Groups_ (1, 14) = 17.58 p=0.0009, F_Hemisphere_ (1, 14) = 9.68 p=0.0077; contra vs ipsi 6-OHDA p=0.0028; 6-OHDA vs CNT contra p=0.0004. **k** In S1BF. Two-way ANOVA with Sidak’s multiple comparisons test F_GroupsXHemisphere_ (1, 14) = 2.08 p=0.17). In blue, CNT n=5 and in orange, 6-OHDA n=4. **l** IMARIS semi-automatic three-dimensional reconstruction and quantitative morphometric analysis of PV-IN. Scale bar, 20 µm. **m** Sholl analysis to study PV-IN morphology complexity in M1. Two-way ANOVA with Sidak’s multiple comparisons test F_Radius_ (139, 1820) = 199.2 p<0.0001, F_Group_ (3, 1820) = 36.83 p<0.0001. **n** Interhemispheric analysis of PV-IN total branching points in control and 6-OHDA mice in M1. Two-way ANOVA with Sidak’s multiple comparisons test F_GroupsXHemisphere_ (1, 14) = 11.82 p=0.0040, F_Hemisphere_ (1, 14) = 11,45 p=0.0045; contra vs ipsi 6-OHDA ***p=0.0009; CNT vs 6-OHDA ipsi *p=0,032. In blue, CNT n=5 and in orange, 6-OHDA n=4. Mean ± SEM. *p≤0.05; **p<0.01; ***p<0.001; ****p<0.0001. See also Supplementary Table 3 and 4.

The density of cortical PV-INs showed a significant decrease in the 6-OHDA group compared to controls. This reduction was consistent across all cortical regions investigated, including M2 (Fig. 3b), M1 (Fig. 3c), and S1BF (Fig. 3d). In M2, only the three most superficial cortical bins were analyzed due to anatomical constraints, yet the results mirrored those seen in the other regions. These observations align with earlier findings from wide-field imaging, suggesting that PV-IN loss is associated with disrupted cortical networks.

The reduction in PV-IN density exhibited a symmetric distribution between hemispheres and consistent uniformity across cortical layers. Interestingly, quantitative analysis of presynaptic puncta-rings encircling non-PV neurons (Fig. 3e) revealed significant differences between experimental groups. Across all the three cortical areas investigated, a marked reduction in punta-ring intensity around non-PV neuronal somata was observed in the contralateral hemispheres of 6-OHDA mice (Fig. 3f-h). Conversely, the ipsilateral hemisphere of 6-OHDA mice showed no significant differences compared to the control and, in M1, presynaptic punta-ring density even appeared to be higher (Fig. 3f-h). These results revealed a discrepancy: despite the loss of PV-INs, inhibitory synaptic contacts did not appear to decrease proportionally. To further investigate this hypothesis, we analyzed the relationship between PV-IN reduction and puncta-ring density by normalizing PV puncta density to PV-IN density. In M2 (Fig. 3i), there was a symmetric increase in the PV puncta density/PV-IN density ratio for 6-OHDA mice by suggesting a heavy compensatory mechanism in both the hemispheres. In M1 (Fig. 3j), the ipsilateral hemisphere of 6-OHDA mice exhibited the most significant increase in the ratio. In contrast, contralateral hemisphere values remained consistent with control levels, by suggesting a lack of compensatory upregulation. In S1BF of 6-OHDA mice (Fig. 3k) instead there was a trend toward a higher ratio in the ipsilateral hemisphere. However, this compensation appears less pronounced compared to the motor regions.

Our results suggest that, despite the symmetric loss of PV-INs, their synaptic output is selectively upregulated in the ipsilateral hemisphere. This mismatch indicates the presence of a compensatory mechanism, potentially involving an increase in the strength of synaptic contacts formed by the long-range PV-INs to counterbalance the loss of inhibitory neurons.

To assess morphological complexity of PV-IN, we performed 3D dendritic reconstructions (Fig. 3l) and 3D Sholl analysis (Fig. 3m), revealing a significant reduction in total branching points in the ipsilateral hemisphere of 6-OHDA-lesioned mice compared to the contralateral side (Fig. 3n). These findings suggest that dopamine depletion may induce dendritic atrophy in PV-INs, potentially disrupting local circuit organization, impairing their ability to maintain inhibitory network stability and recruiting long-range connections.

### PV-INs Functional Connectivity Predicts Pathology Progression

We then aimed to assess the potential of FC as a biomarker for disease progression in the 6-OHDA model of PD. To achieve this, we first integrated the entire dataset, encompassing both physiological and histological data, and performed principal component analysis (PCA) to determine whether the two groups could be distinctly separated. Indeed, PCA revealed clear clustering of control (CNT) and 6-OHDA groups along the first principal component (PC1), accounting for 46.5% of the variance (Fig. 4a). Among all features contributing to PC1, FC data ranked prominently, alongside puncta density and lesion assessments in the CPu, underscoring FC’s relevance as a descriptor of disease progression (Fig. 4b). To further validate our findings, we trained a support vector machine (SVM) classifier using the entire dataset and various subsets. The classifier achieved an impressive 89% accuracy when trained with FC data from wide field recordings, while performance dropped to chance level (50%) in the shuffled (bootstrapped) condition, where data was randomly assigned to animals (Fig. 4c). We then examined patterns of correlations within the entire dataset using a correlation matrix (Fig. 4d). This analysis highlighted both positive and negative associations among the measured variables, with notable strong correlations observed between behavioral measures, FC parameters, and synaptic markers. Interestingly, much of the correlation between FC and other parameters was significant for FC data from the right hemisphere, corresponding to the injection site. An intriguing exception to this pattern was observed in the retrosplenial cortex, warranting further investigation (Fig. 4d). Examining specific correlations, we found that WF connectivity in cortical areas was strongly associated with the severity of motor deficits, underscoring its potential as a biomarker for disease progression. This relationship was particularly evident in cortical regions critical for motor processing, where alterations in PV-IN connectivity closely mirrored behavioral impairments (Fig. 4e). Moreover, PV-IN density exhibited a robust correlation with WF connectivity, suggesting that structural changes in inhibitory networks are intrinsically linked to functional connectivity deficits (Fig. 4f). Collectively, these findings indicate that while PV-IN density decreases across hypoconnected cortical regions in the 6-OHDA model, the concurrent reorganization of PV-IN synaptic connectivity—especially in the ipsilateral hemisphere— may represent a compensatory mechanism. This reorganization provides valuable insight into cortical dysfunction in PD models by highlighting the complex interplay between neuronal density and synaptic adaptations in preserving cortical network stability.

**Fig. 4.**
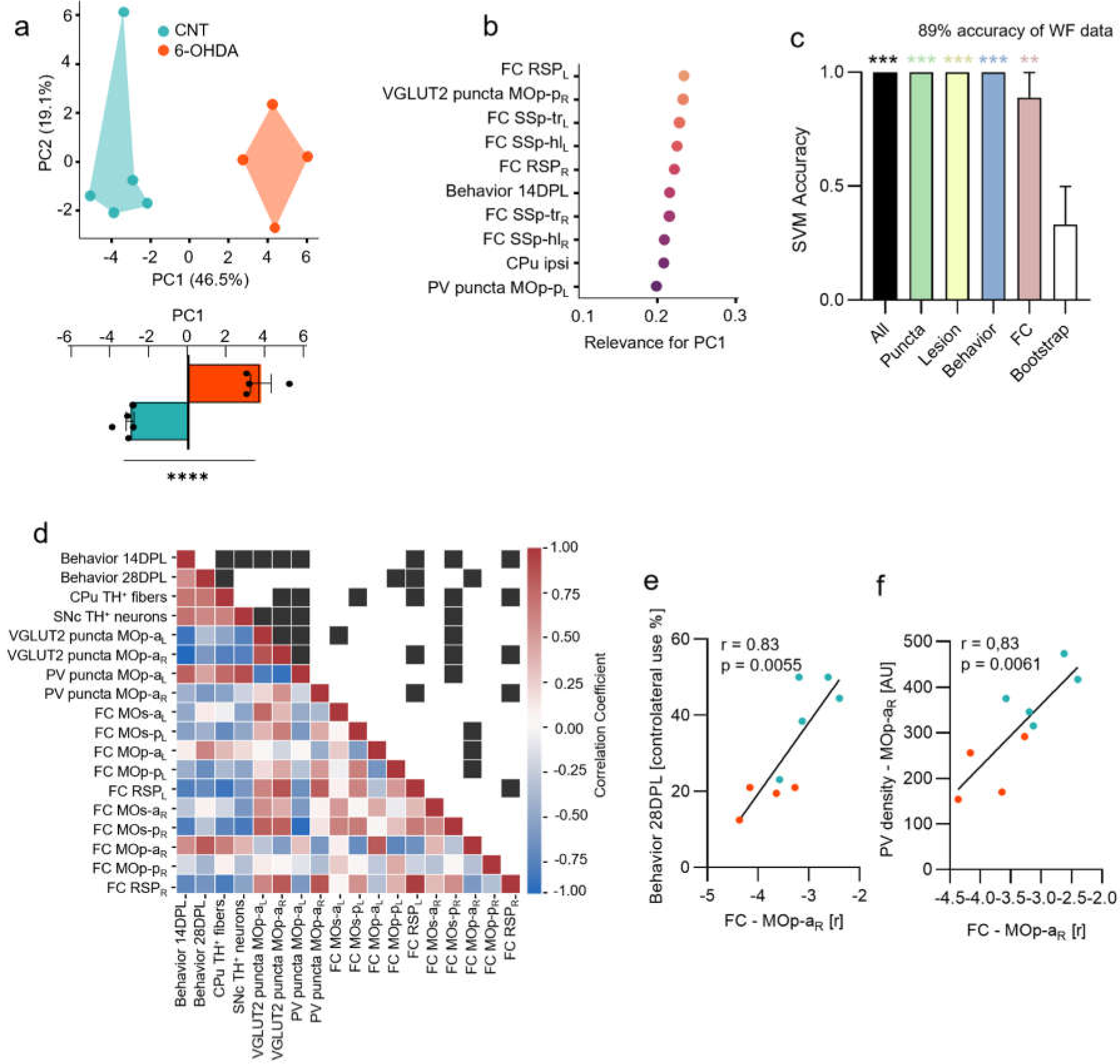
PV-INs Functional Connectivity Predicts Pathology Progression. **a** Scatter plot on the Principal Component space showing the distribution of CNT and 6-OHDA mice on the first and second components. Unpaired t-test p<0.0001. **b** First ten ranked features contributing to the PC1. **c** Average classification accuracy of the SVM classifier in distinguishing CNT and 6-OHDA mice using Leave-One-Out Cross-Validation (LOO-CV). The SVM was trained on different subsets of the dataset plus a bootstrap dataset as control, and accuracy was measured across all Leave-One-Out splits (Wilcoxon Signed Rank Test (against chance: 0.5): All, p=0.0039; Puncta, p=0.0039; Lesion, p=0.0039; Behavior, p=0.0039; WF, p=0.0391). **d** Heat map representing values from the correlation matrix (Pearson correlation) of analysed parameters (bottom triangle), and the significance of each pairwise comparison (top triangle: black box indicates p < 0.05). **e-f** Scatter plot and linear regression detailing specific correlations depicted in **D**. In particular, we show the correlation between wide-field data at 28DPL in the MOp-a_R_ with **e** performance in the behavioral test at 28DPL, and **f** PV density in MOp-pR (r and p-values are indicated in the figures). All the FC data refer to FC at 28DPL. In blue, CNT n=5 and in orange, 6-OHDA n=4. Mean ± SEM. *p≤0.05; **p<0.01; ***p<0.001; ****p<0.0001

### Excitatory/Inhibitory Imbalance and Microglial Response in the Motor Cortex Following Dopaminergic Neurodegeneration

The striatum communicates indirectly with the motor cortex via basal ganglia circuitry while also receiving direct cortical modulation (Supplementary Fig. 4a). Given our observation of alterations in PV-IN connectivity—crucial for maintaining synchronized cortical activity—we investigated whether dopaminergic loss induces homeostatic changes in excitatory/inhibitory (E/I) balance within the motor cortex. To assess synaptic integrity, we analyzed cortical vesicular markers for both inhibitory (VGAT, Fig. 5a) and excitatory (VGLUT1, Fig. 5c; VGLUT2, Fig. 5e) neurotransmission. Specifically, we quantified the mean fluorescence of excitatory and inhibitory terminals impinging on the soma of layer II/III neurons and further examined synaptic integrity by evaluating vesicular markers as puncta-rings surrounding the cell body. Our findings revealed a pronounced E/I imbalance in the cortex of 6-OHDA-lesioned mice, characterized by significant alterations in the density of these synaptic markers, highlighting potential compensatory mechanisms in response to dopaminergic degeneration.

**Fig. 5.**
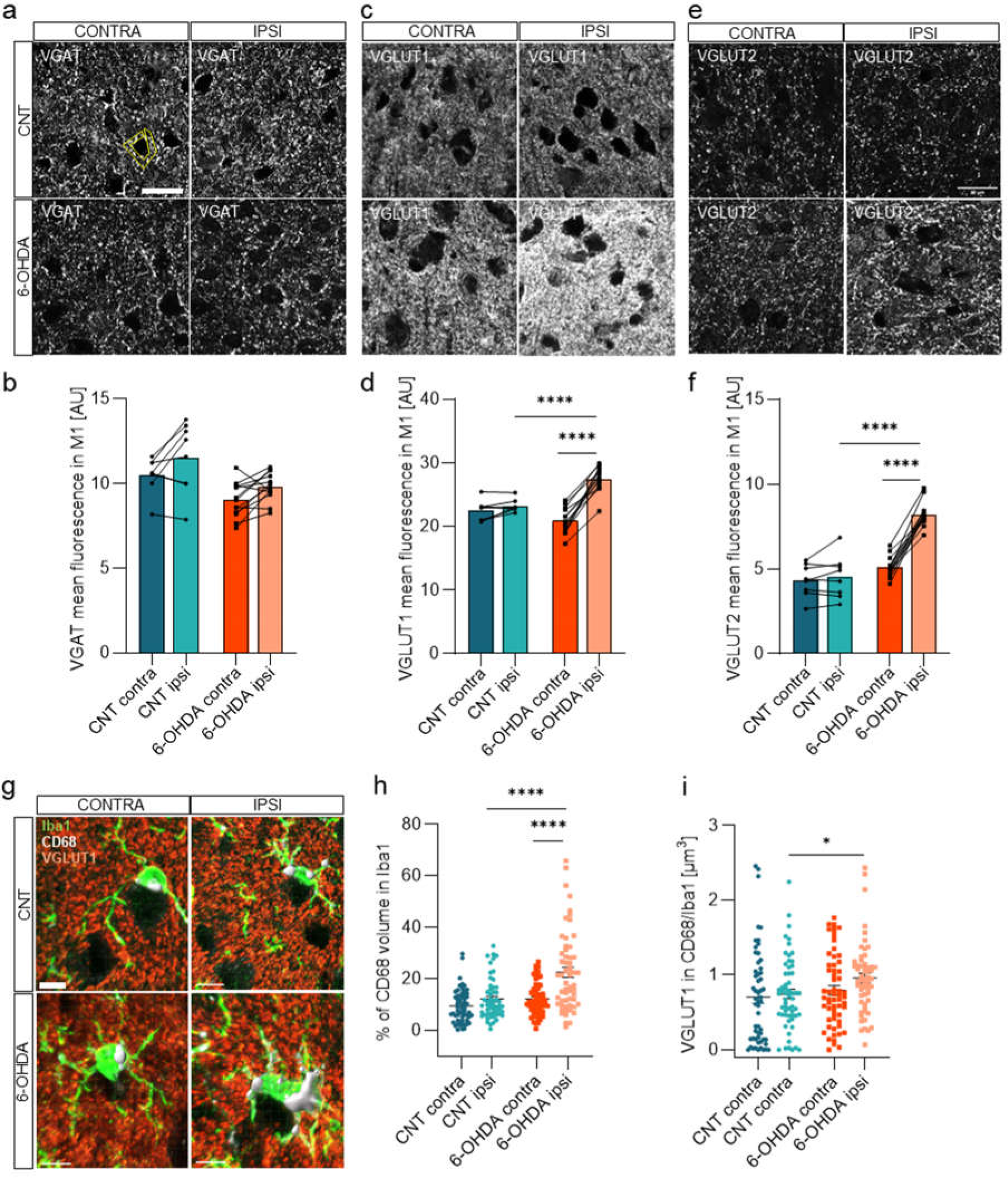
Modulation of excitatory and inhibitory transporters and microglia activation after 6-OHDA injection. **a** VGAT puncta-rings analysis. Magnified view of the primary motor cortex stained with anti-VGAT for control (in blue, CNT n=8) and 6-OHDA (in orange, 6-OHDA n=13) groups. Scale bar: 20 µm. **b** Interhemispheric analysis of anti-VGAT fluorescence mean values of puncta-rings around cell bodies per animal in control and 6-OHDA group, calculated in mean fluorescence of puncta-rings around cell bodies. Data from three fields of view per three brain sections n=72 cells for each dataset and CNT animals n=8 vs 6-OHDA animals n=13 were analysed. Two-way ANOVA with Sidak’s multiple comparisons test F_GroupsXHemisphere_ (1, 38) = 0.01178 p=0.9144, F_Groups_ (1, 38) = 10.58 p=0.0024, F_Hemisphere_ (1, 38) = 3.048 p=0.0.0889. **c VGLUT1 puncta-rings analysis in contro**l (in blue, CNT n=8) and 6-OHDA (in orange, 6-OHDA n=13) groups. Magnified view of the primary motor cortex stained with anti-VGLUT1. Scale bar: 20 µm. **d** Interhemispheric analysis of anti-VGLUT1 fluorescence mean values of puncta-rings in control and 6-OHDA group. Data from three fields of view per three brain sections n=72 cells for each dataset and CNT animals n=8 vs 6-OHDA animals n=13 were analysed. Two-way ANOVA with Sidak’s multiple comparisons test F_GroupsXHemisphere_ (1, 38) = 26.21 p<0.0001, F_Groups_ (1, 38) = 5.374 p=0.026, F_Hemisphere_ (1, 38) = 40.01 p<0.0001; ipsi 6-OHDA vs CNT p<0.0001; 6-OHDA vs CNT ipsi p<0.0001. **e** VGLUT2 puncta-rings analysis in control (in blue, CNT n=8) and 6-OHDA (in orange, 6-OHDA n=13) groups. Magnified view of the primary motor cortex stained with anti-VGLUT2. Scale bar: 20 µm. **f** Interhemispheric analysis of anti-VGLUT2 fluorescence mean values of puncta-rings in control and 6-OHDA group. Data from three fields of view per three brain sections n=72 cells for each dataset and CNT animals n=8 vs 6-OHDA animals n=13 were analysed. Two-way ANOVA with Sidak’s multiple comparisons test F_GroupsXHemisphere_ (1, 38) = 27.21 p<0.0001, F_Groups_ (1, 38) = 64.94 p<0.0001, F_Hemisphere_ (1, 38) = 36.56 p<0.0001; contra vs ipsi 6-OHDA p<0.0001; 6-OHDA vs CNT ipsi p<0.0001. **g-i** Phagocytosis analysis of Iba1-positive cells in the primary motor cortex. **g** Representative tridimensional surface analysis of Iba1, CD68 and VGLUT1 markers in the primary motor cortex stained with anti-Iba1, anti-CD68 and anti-VGLUT1. Scale bar: 7 µm. **h** Interhemispheric analysis of phagosome volume (CD68+) inside Iba1+ cells. n=60 cells for each dataset and CNT animals n=4 vs 6-OHDA animals n=5 were analysed. Two-way ANOVA with Sidak’s multiple comparisons test F_GroupsXHemisphere_ (1, 228) = 9.03 p=0.0030, F_Groups_ (1, 228) = 25.85 p<0.0001, F_Hemisphere_ (1, 228) = 26.08 p<0.0001; contra vs ipsi 6-OHDA p<0.0001; 6-OHDA vs CNT ipsi p<0.0001. **i** Interhemispheric analysis of VGLUT1 engulfment inside CD68 volume, normalized on the total amount of VGLUT1 in the field. n=60 cells for each dataset and CNT animals n=4 vs 5 6-OHDA animals n=5 were analysed. Two-way ANOVA with Sidak’s multiple comparisons test F_Groups_ (1, 228) = 5.00 p=0.026; 6-OHDA vs CNT ipsi p=0.053. Mean ± SEM. *p≤0.05; **p<0.01; ***p<0.001; ****p<0.0001. See also Supplementary Figure 4 and Supplementary Table 5.

For VGAT, expression levels remained stable, with no significant changes observed, suggesting that inhibitory neurotransmission, represented by VGAT puncta-ring density, was unaffected by dopaminergic loss at the group level. Interhemispheric analysis of anti-VGAT fluorescence mean values confirmed this, showing no significant interaction between the animal groups and hemispheres (Fig. 5b). In contrast, for VGLUT1, a significant increase was observed in the ipsilateral hemisphere, suggesting that excitatory synaptic transmission was globally enhanced following dopaminergic lesion. The interhemispheric comparison further reinforced this trend, with pronounced differences observed in the ipsilateral hemisphere of the 6-OHDA group compared to controls (Fig. 5d). Similarly, VGLUT2 expression was significantly upregulated in ipsilateral (Fig. 5f, highlighting that the E/I imbalance in the 6-OHDA model is largely driven by elevated mean fluorescence of puncta-rings in the ipsilateral hemisphere. These findings suggest an adaptive response to the loss of dopaminergic input, characterized by a pronounced E/I imbalance, likely aimed at compensating for the reduced inhibitory influence in the injured cortex.

These findings suggest that the loss of dopaminergic input triggers an adaptive response in the motor cortex, characterized by a pronounced E/I imbalance, likely aimed at compensating for the reduced dopaminergic and altered PV-IN influence in the injured hemisphere. Given the crucial role of microglial in synaptic remodeling, we further examined their involvement in these cortical alterations. As the brain’s resident phagocytes, microglia actively remove dead and dying neurons, as well as synapses [31]. In PD and related models like the 6-OHDA lesion, microglial activation often parallels dopaminergic neuron degeneration and is thought to exacerbate or even trigger pathological changes in the cortex [32, 33]. Given microglia’s pivotal role in synaptic remodeling especially during brain injury [34–36], we thus investigated their role in the observed synaptic modulation in 6-OHDA treated mice. First, we analysed microglial density in cortical areas with identified connectivity deficits and observed no significant changes in the primary motor cortex (M1, Supplementary Fig. 4b), secondary motor cortex (M2, Supplementary Fig. 4c), and somatosensory barrel field (S1BF, Supplementary Fig. 4d).

Then, we proceeded to quantify microglial phagocytic ability using CD68 as a marker (Fig. 5g). CD68 is a lysosomal and endosomal marker associated with phagocytosis, specifically expressed by macrophages, including Iba1-positive microglia in the CNS. [37] Interestingly, we found an increase in CD68 in microglia in the ipsilateral cortex of 6-OHDA treated mice, suggesting an enhanced microglia engagement in phagocytic activity (Fig. 5h). Additionally, there was an increased engulfment of VGLUT1 by microglia (Fig. 5i). This suggests that microglia are actively participating in the removal of synaptic components, which may be a response to neuronal damage and synaptic remodeling associated with dopaminergic degeneration.

## DISCUSSION

This study provides compelling evidence that subcortical dopaminergic degeneration in a 6-OHDA induced Parkinson’s disease model [1, 38], significantly impacts not only cortico-striatal electrophysiological coupling but also functional and anatomical cortical features related to the inhibitory system, specifically Parvalbumin-positive interneurons (PV-INs). Through a combination of electrophysiological recordings, wide-field calcium imaging, and histological analyses, we uncovered a remodelling mechanism of cortical plasticity involving PV-INs in response to nigrostriatal degeneration. Electrophysiological analysis of both the striatum and motor cortex, focusing on their functional coupling in resting state, revealed a significant increase in striatal delta band activity in 6-OHDA mice, consistent with previous studies (Fig. 1c) [39, 40]. Although direct changes in motor cortex activity at rest were not observed, we identified a progressive increase in cortico-striatal coherence in the delta band as the pathology advanced (Fig. 1f). Interestingly, the rise in striatal delta power alone does not fully account for the observed increased cortico-striatal coherence, as these phenomena can occur independently though related [41]. In fact, elevated striatal delta power reflects increased local oscillatory activity, whereas increased coherence indicates stronger functional connectivity or phase alignment between the cortex and striatum. This distinction suggests that the 6-OHDA-induced neurodegeneration drives pathological entrainment of cortico-striatal communication, potentially contributing to disrupted cortical network dynamics in Parkinson’s disease.

During voluntary movement, we instead observed stark differences in spectral modulation between 6-OHDA-treated and control mice. In the striatum of 6-OHDA mice, spectral modulation was essentially absent from the earliest recording stages, indicating that the neurotoxin rapidly disrupts movement-related subcortical oscillatory activity (Fig. 1i). In contrast, control mice exhibited robust movement-related spectral modulation in the striatum during the voluntary retraction task, underscoring the early impact of 6-OHDA on striatal function (Fig. 1j).

Interestingly, in control mice, spectral modulation gradually decreased in both the striatum and cortex over the recording weeks (Fig. 1i, j). This decline may reflect two factors: a potential loss of electrode signal quality over time, as is common in chronic recordings [42], and a possible habituation effect to the task, where repeated exposure reduces the neuronal response [43]. Despite this overall trend, we observed an opposing pattern in the high-gamma range in the motor cortex of 6-OHDA mice, where high-gamma power progressively increased throughout recording weeks (Fig. 1o). This heightened high-gamma modulation may reflect network hypersynchrony or enhanced firing near the electrode tip [44], potentially linked to reduced parvalbumin-positive interneurons (PV-INs) signaling. These findings underscore the dual role of cortical and subcortical dynamics in PD pathology, where the motor cortex undergoes spectral changes to compensate for striatal dysfunction. However, the progressive divergence in spectral patterns, particularly the rise in high-gamma modulation during movement, highlights the maladaptive nature of these compensations.

Previous research showed that PV-INs are key regulators of cortical inhibition control and suggested that gamma oscillations regulated by PV-INs promote synaptic plasticity in cortical motor areas [7, 11, 29, 45–47]. We therefore longitudinally investigated alterations in PV-INs cortical network using wide-field calcium imaging (Fig. 2). Electrophysiological and imaging analyses revealed that while the pathological phenotype remains stable over time (Supplementary Fig. 1f), FC undergoes progressive deterioration with the most pronounced alterations emerging at 28 days post-lesion (Fig. 2f). Critically, inter-hemispheric connectivity was strongly affected already at early time points (Fig. 2h), indicating that dopaminergic loss impacts cortical communication between hemispheres more than intra-hemispheric connections. Although our findings suggest a disruption of inter-hemispheric communication, further investigation into PV-IN density, morphology and synaptic connectivity (Fig. 3) provides critical insights into the role of the healthy hemisphere in the compensatory mechanisms that reshape PV-IN cortical network. Despite a symmetric reduction in PV-IN density across hemispheres and cortical layers (Fig. 3b-d), we observed a selective upregulation of inhibitory synaptic output in the ipsilateral hemisphere to the lesion, particularly strong in the primary and secondary motor cortex (Fig. 3i-k). At the same time, 3D dendritic reconstructions and Sholl analysis of PV-INs in the primary motor cortex revealed a significant reduction in dendritic branching in layers 2/3 PV-INs of the ipsilateral hemisphere (Fig. 3n). In contrast, PV-INs in the contralateral hemisphere exhibit increased dendritic branch complexity (Fig. 3n). These findings suggest a shift in inter-hemispheric control, with the contralateral hemisphere compensating for functional deficits by enhancing long-range connectivity and synaptic contacts.

PV-INs in cortical layers 2/3 possess relatively long axons that contact excitatory (pyramidal) and PV+ cells, facilitating the synchronization of gamma oscillations[12]. These synchronized oscillations are integral to motor planning and execution and suggest a potential indirect influence on interhemispheric communication through long-range pyramidal neurons, particularly in layers 2/3. [48] Such changes in PV-IN distribution and synaptic activity highlight a broader disruption in the inhibitory network, extending beyond PV-IN cell bodies to their synaptic targets. This disruption likely compromises inhibitory control mechanisms, impairing the generation and maintenance of gamma oscillations critical for cortical network stability. These structural alterations align with functional connectivity deficits, emphasizing the essential role of PV-IN integrity in maintaining cortical stability. Moreover, our findings contribute to a broader understanding of cortical dysfunction in PD models by highlighting the intricate relationship between neuronal density and synaptic adjustments in maintaining cortical network stability.

Overall, our results suggest that dopaminergic loss does not lead solely to a breakdown of PV-INs cortico-cortical communication but instead triggers an asymmetric gain-of-function adaptation in the contralateral hemisphere, likely as a mechanism to offset motor and sensory deficits.

Atrophy of dendrite branching points in the ipsilateral hemisphere of 6-OHDA mice and the structural change following degeneration of the dopaminergic system and consequent decrease of subcortical innervation to motor cortex is compensated with an enhanced synaptic strength. Globally, the observed symmetric reductions in PV cell bodies could reflect a widespread vulnerability of these neurons. Locally, in the ipsilateral hemisphere, the reduced connectivity among PV-PV circuits and the decrease in dendritic complexity may limit PV-IN functionality. This could necessitate compensatory synaptic remodeling, such as increased PV-to-non-PV connections. These changes might contribute to cortical network instability and altered oscillatory activity, potentially exacerbating motor deficits characteristic of Parkinson’s disease.

The observed changes are consistent with previous reports of PV-IN vulnerability in neurodegenerative conditions [49, 50] and underscore their importance as therapeutic targets in PD. The loss of PV-INs, which play a central role in maintaining excitatory/inhibitory balance, may exacerbate cortical hyperexcitability and contribute to the E/I imbalance observed in this study. By impairing gamma oscillations and long-range synchrony, PV-IN reductions likely disrupt motor planning and execution, key features of PD pathology. Future therapeutic strategies should focus on restoring PV-IN function or compensating for their loss to stabilize cortical networks and improve motor outcomes in PD. Indeed, histological analysis revealed a profound imbalance between excitatory and inhibitory synaptic markers [51, 52]. with increased excitatory drive likely compensating for dopaminergic loss in primary motor cortex. While VGAT levels (inhibitory synaptic marker) remained stable, VGLUT1 and VGLUT2 (excitatory synaptic markers) were significantly upregulated. This imbalance exacerbates cortical hyperactivity and disrupts normal motor processing. This suggests that cortical hyperactivity emerges as a compensatory response to subcortical degeneration. Interestingly, this excitatory/inhibitory imbalance cannot be attributed to dopaminergic degeneration alone, as the motor cortex does not receive direct dopaminergic innervation [53, 54]. Instead, we propose glutamatergic modulation plays a critical role, as the primary motor cortex integrates subcortical inputs and modulates striatal dopaminergic neurons [55–57].

In addition to neuronal and synaptic changes, we observed significant alterations in microglial activity in the motor cortex. Microglial cells displayed increased phagocytic activity, suggesting an ongoing neuroinflammatory response. Interestingly, we found an increase in VGLUT1 phagocytosis by microglia, which is captivating since total VGLUT1 in the tissue seems to be elevated. This apparent paradox may be explained by a compensatory upregulation of excitatory synapse formation in response to increased microglial activity, suggesting that neurons might bolster synaptic connectivity to offset potential losses. Alternatively, the accumulation of VGLUT1 could indicate that engulfed material is not fully degraded, thereby contributing to the overall increase. These findings underscore a dynamic interplay between synaptic remodeling and microglial clearance, hinting that heightened synaptic turnover may underlie the observed phenomena. Such a scenario suggests a dynamic interplay between compensatory synaptic remodeling and neuroinflammatory processes, exacerbating neuronal damage while attempting to preserve cortical functionality [58–60]. While this response may initially aim to clear debris and maintain homeostasis, excessive microglial activation could exacerbate cortical dysfunction and contribute to disease progression. This evidence underscores the need to address both neuronal and microglial contributions in future therapeutic strategies.

Collectively, our findings point to a cascade of maladaptive cortical processes initiated by subcortical dopaminergic degeneration. High-gamma cortical hypersynchrony and pathological striatal delta modulation disrupt cortico-striatal communication, impairing motor control. Reduced PV-IN density and connectivity destabilize cortical networks, leading to hyperexcitability and compensatory plasticity, with hyper-synchronized activity and a possible overcoming of the contralateral hemisphere. PV alterations lead to E/I imbalance and microglial phagocytosis amplifies synaptic turnover, linking neuroinflammation to synaptic modelling and further circuit dysfunction. This cascade underscores the complex interplay between neuronal, synaptic, and inflammatory mechanisms in PD pathology. Taken together, these disruptions in local and global network dynamics contribute to motor deficits in PD. The progressive nature of these alterations emphasizes the importance of early intervention to prevent or mitigate cortical dysfunction.

Importantly, this study identifies the motor cortex as a critical site for therapeutic intervention in PD. Our results suggest that wide-field calcium imaging could serve as a valuable biomarker for monitoring disease progression and evaluating therapeutic efficacy. The observed functional reorganization and both electrophysiological and synaptic imbalances provide critical insights into cortical compensatory mechanisms, reinforcing the need for therapeutic strategies targeting the motor cortex. Non-invasive approaches, such as transcranial alternating current stimulation, might offer promise for modulating abnormal cortical activity and synchronizing cortico-striatal circuits [61]. Additionally, altered microglial activity suggests that neurotrophic factors, such as nerve growth factor (NGF), may help restore neuronal function and mitigate neuroinflammation [62]. In conclusion, addressing motor cortex dysfunction is essential for slowing PD progression and improving motor function. A combination of approaches, such as brain stimulation to correct electrophysiological abnormalities and NGF to support neuronal survival, might offer a synergistic path forward.

### Limitations of the study

While this study provides valuable insights into the 6-OHDA mouse model of Parkinson’s disease, several limitations must be acknowledged. The 6-OHDA model effectively replicates key aspects of dopaminergic neuron loss; however, it might not fully encompass the progressive and bilateral nature of human PD. In this study, we opted for a unilateral lesion approach to establish a well-characterized model of the pathology. Nevertheless, exploring bilateral lesion in future studies could be useful to further assess therapeutic efficacy in a more widespread neurodegenerative context, improving translational relevance to clinical settings.

Additionally, this study primarily focuses on motor deficits, while non-motor symptoms, such as cognitive and emotional dysfunctions, remain unexplored. Since our objective was to investigate the motor cortex as a potential therapeutic target, we employed a model that effectively recapitulates motor symptoms. However, incorporating assessments of non-motor dysfunctions in future studies might provide a more comprehensive understanding of the disease.

Furthermore, longer-term studies could be valuable to fully understand the chronic effects of dopaminergic degeneration. Another consideration is the interpretation of cortico-striatal dynamics, as this study does not allow for a direct causal relationship between the cortex and striatum to be established. This suggests the need for further exploration of multisynaptic pathways or shared synaptic drives that might explain these findings.

Finally, although the results highlight potential therapeutic targets, such as non-invasive brain stimulation and nerve growth factor (NGF), further investigations in more complex models or clinical settings would be necessary to validate these interventions before they can be considered for translation to human therapies.

### Conclusions

Our study highlights the critical role of cortical PV-INs, oscillatory dynamics, and neuroinflammatory processes in PD. The findings reveal that delta and gamma oscillations represent key markers of cortico-striatal dysfunction. PV-INs serve as central regulators of cortical stability, with their dysfunction exacerbating E/I imbalance and motor impairments. Synaptic remodeling and microglial activity contribute to the pathological adaptations, linking neuroinflammation to neuronal damage. These insights offer a framework for understanding PD-related cortical changes and suggest potential therapeutic targets. Interventions aimed at restoring PV-IN functionality, modulating oscillatory activity, or mitigating neuroinflammation could provide novel strategies to alleviate motor and cognitive symptoms in PD. Future research should explore the temporal dynamics of these processes and their interactions to develop targeted therapies aimed at improving clinical outcomes and slowing disease progression.

## FUNDING

Fondo Beneficenza Intesa San Paolo n. B/2022/0193, Project “ONDA”, Project Manager: Cristina Spalletti

## RESOURCE AVAILABILITY

### Lead contact

Further information and requests for resources and reagents should be directed to and will be fulfilled by the lead contact, Cristina Spalletti.

### Materials availability

All source data are accessible in a public database.

## ACKNOWLEDGMENTS

We thank Francesca Biondi (CNR Pisa) for the excellent animal care, Maria Pasquini, Elena Novelli and Renzo di Renzo for technical support with imaging and informatics.

## AUTHOR CONTRIBUTIONS

Conceptualization, C.S., A.L.A.M., A.Maz., and S.C.; methodology, C.S., A.L.A.M., A.Maz., and S.C.; investigation, A.Mi., F.M., A.Mar., E.M., E.C., A.T., N.M., A.Maz., A.L.A.M., C.S., and S.C.; writing – original draft, A.Mi., E.M., N.M., A.Maz., A.L.A.M., S.C., and C.S., writing – review & editing: A.Mi., E.M., E.C., A.T., N.M., A.Maz., A.L.A.M., S.C., and C.S; funding acquisition, C.S., A.L.A.M., and A.Maz.; resources, C.S., A.L.A.M., and A.Maz., supervision, C.S., A.L.A.M., A.Maz., and S.C.

## DECLARATION OF INTERESTS

Authors declare that they have no competing interests.

## SUPPLEMENTAL INFORMATION

**Document S1. Figures S1–S4 and Table S1-S5**.

## Notes

### Competing Interest Statement

The authors have declared no competing interest.

