## Supplementary material for "Altered cortical network in Parkinson’s Disease: the central role of PV interneuron and synaptic remodelling"

### Supplementary Information

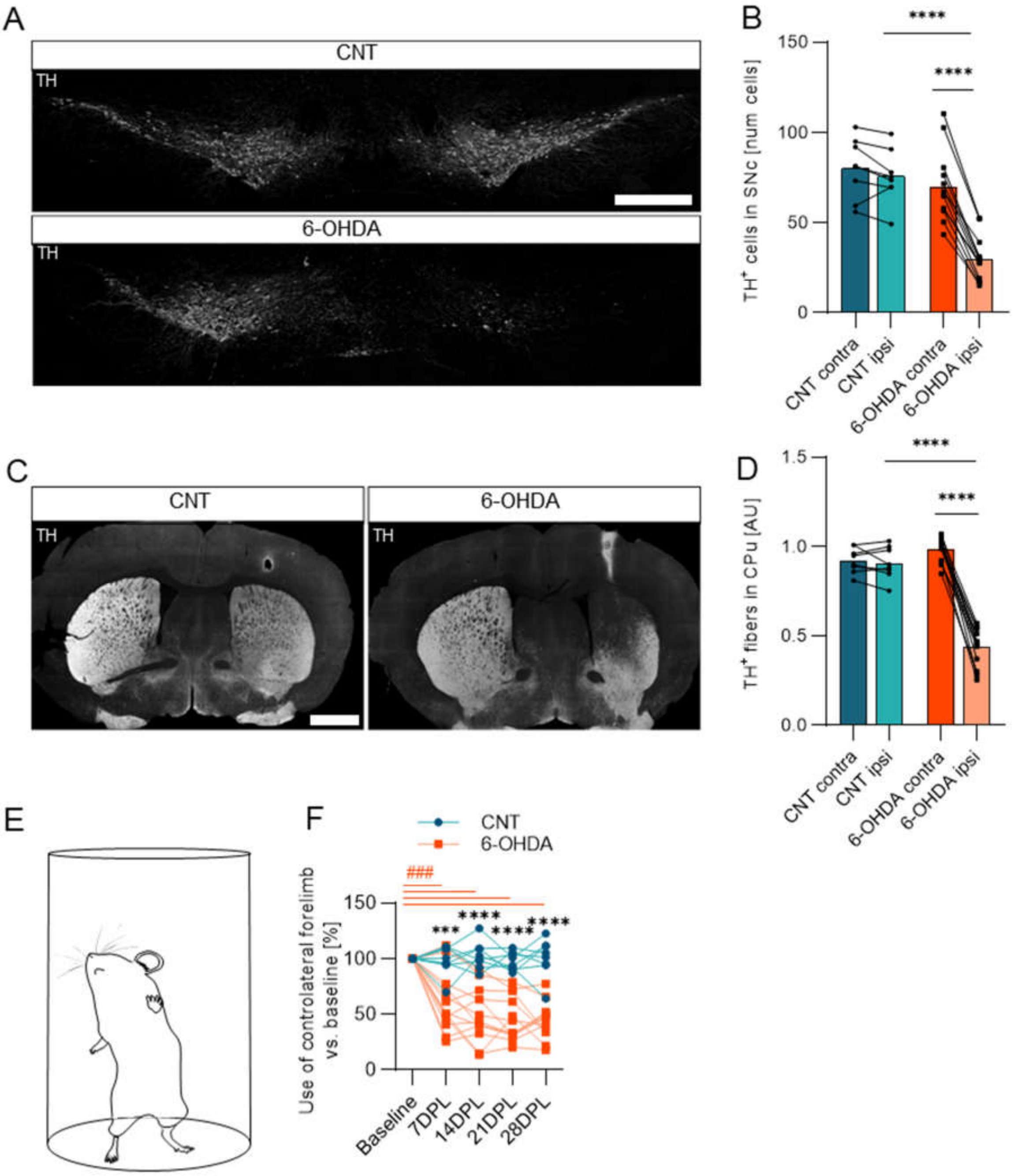


**Figure S1.**

**(A)** Representative coronal sections of the SNpc stained for TH in CNT (top) and 6-OHDA mice (down). Scale bar, 500 µm.

**(B) Quantification of TH-positive neurons in SNc.** Quantification of $TH^+$ neurons in both contralateral (dark color) and ipsilateral (light color) hemisphere to the lesion. CNT group (contra = 80.2 ± 5.9; CNT ipsi = 75.7 ± 5.3). 6-OHDA group (contra = 69.9 ± 5.4; ipsi = 29.7 ± 3.4). Two-way ANOVA with Sidak's multiple comparisons test $F_{GroupsXHemisphere}$ (1, 38) = 12.33 p=0.0012, $F_{Groups}$ (1, 38) = 30.61 p<0.0001, $F_{Hemisphere}$ (1, 38) = 19.24 p<0.0001; contra vs ipsi 6-OHDA p<0.0001; 6-OHDA vs CNT ipsi p<0.0001).

**(C)** Representative sections of striatum stained for TH from control (left) and 6-OHDA lesioned mice (right). Scale bar, 1 mm.

**(D) Quantification of TH-positive fibers in CPu.** Quantification of average relative optical density (ROD) for $TH^+$ fibers fluorescence in both contralateral (light color) and ipsilateral (dark color) hemisphere to the lesion. CNT group (contra = 0.92 ± 0.03; ipsi = 0.91 ± 0.03). 6-OHDA group (contra = 0.99 ± 0.02; ipsi = 0.44 ± 0.03). Two-way ANOVA with Sidak's multiple comparisons test $F_{GroupsXHemisphere}$ (1, 38) = 92.40 p<0.0001, $F_{Groups}$ (1, 38) = 53.77 p<0.0001, $F_{Hemisphere}$ (1, 38) = 104.0 p<0.0001; contra vs ipsi 6-OHDA p<0.0001; 6-OHDA vs CNT ipsi p<0.0001).

**(E)** Representative schematic representation of Shallert cylinder test.

**(F) Forelimbs use asymmetry in the longitudinal cylinder test.** The percentage of wall contacts made with the contralateral forepaw during cylinder test at 7, 14, 21 and 28 DPL. The CNT group (Baseline = 100 ± 0.0; 7DPL = 98.8 ± 4.2; 14DPL = 102.9 ± 4.5; 21DPL = 98.4 ± 2.8; 28DPL = 102.2 ± 5.0 ). 6-OHDA group (Baseline = 100 ± 0.0; 7DPL = 60.7 ± 7.3; 14DPL = 49.9 ± 6.7; 21DPL = 44.9 ±5.3;

28DPL = 44.5 ± 4.5). Repeated Two-way ANOVA with Sidak's multiple comparisons test $F_{TimeXGroups}$ (4, 76) = 16.88 p<0.0001, $F_{Time}$ (4, 76) = 16.54 p<0.0001, $F_{Groups}$ (1, 19) = 47.46 p<0.0001, $F_{Subjects}$ (19, 76) = 5.143 p<0.0001; 7DPL CNT vs 6-OHDA p<0.0006, 14DPL CNT vs 6-OHDA p<0.0001; 21DPL CNT vs 6-OHDA p<0.0001; 28DPL CNT vs 6-OHDA p<0.0001; One sample t-test with Holm-Sidak's multiple comparisons 6-OHDA 7DPL ###p=0.0004; 6-OHDA 14DPL ###p=0.0004; 6-OHDA 21DPL ###p=0.0004; 6-OHDA 28DPL ###p=0.0004. In blue, CNT n=8 and in orange, 6-OHDA n=13 Data are expressed as mean ± SEM. *p≤0.05; **p<0.01; ***p<0.001; ****p<0.0001

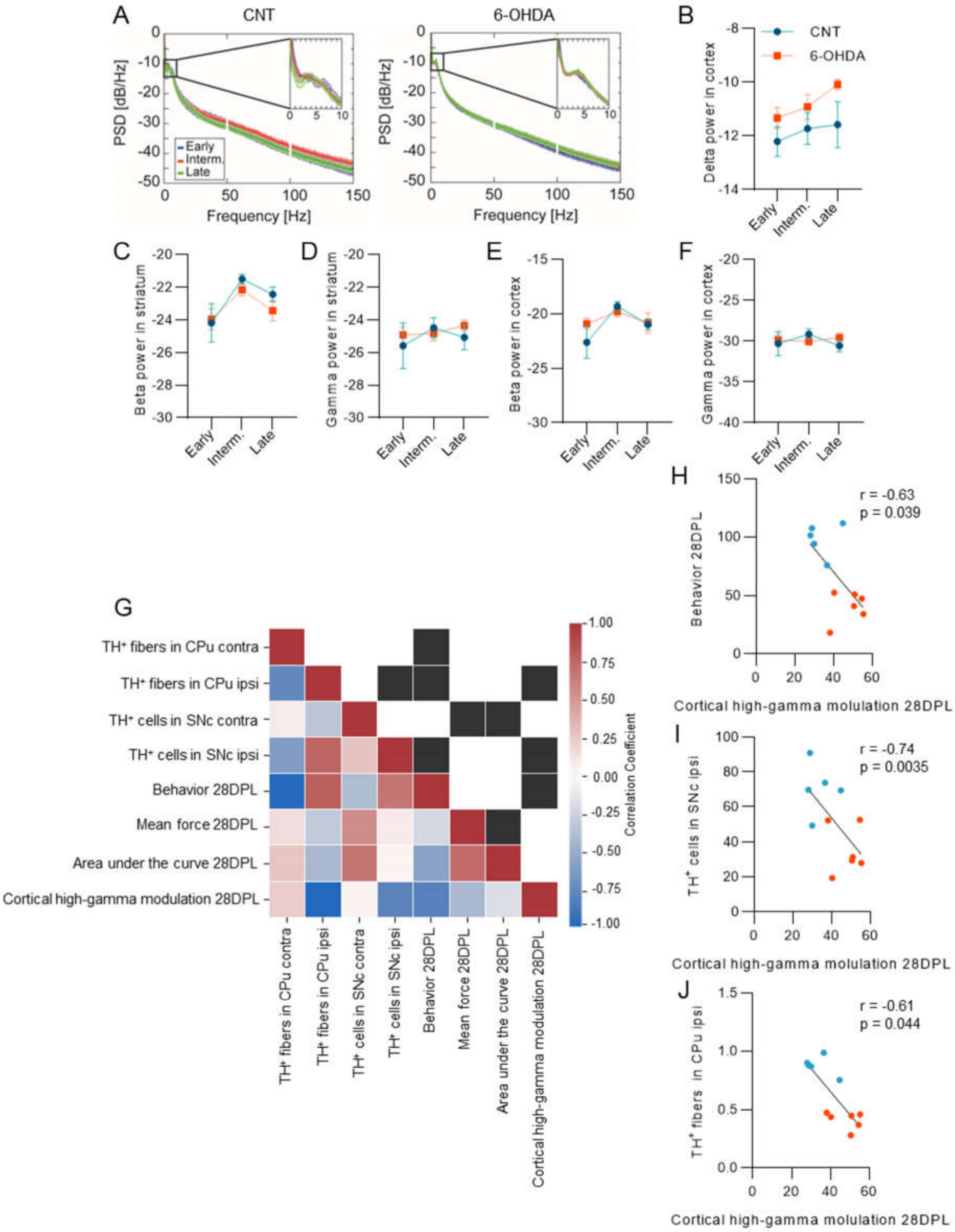


**Figure S2.**

**(A)** Average power spectral density of LFPs in the cortex of the control group (left, n=8) and 6-OHDA mice (right, n=13) across recording weeks. Zoomed inset provides focus on the low-frequencies part of the PSD.

**(B)** Longitudinal cortical power of delta oscillations in controls in blue (CNT n=8) and 6-OHDA mice in orange (n=13). CNT early stage: -12.67 ± 0.73; CNT intermediate stage: -11.98 ± 0.69; CNT late stage: -12.07 ± 1.02; 6-OHDA early stage: -11.33 ± 0.39; 6-OHDA intermediate stage: -10.91 ± 0.46; 6-OHDA late stage: -10.09 ± 0.20. Two-way ANOVA $F_{StagesXGroups}$= 0.28 p=0.75, $F_{Stages}$= 0.83 p=0.44, $F_{Groups}$= 7.22 p=0.008.

**(C)** Longitudinal striatal power of beta oscillations in controls in blue (CNT n=8) and 6-OHDA mice in orange (n=13). CNT early stage: -24.19 ± 1.17; CNT intermediate stage: -21.50 ± 0.30; CNT late stage: -22.44 ± 0.45; 6-OHDA early stage: -23.96 ± 0.64; 6-OHDA intermediate stage: -22.14 ± 0.39; 6-OHDA late stage: -23.44 ± 0.63. Two-way ANOVA with Sidak's multiple comparisons test $F_{StagesXGroups}$ (2, 163) = 0.49 p=0.61, $F_{Stages}$ (2, 163) = 8.12 p=0.0004, $F_{Groups}$ (1, 163) = 0.97 p=0.33.

**(D)** Longitudinal striatal power of gamma oscillations in controls in blue (CNT n=8) and 6-OHDA mice in orange (n=13). CNT early stage: -25.59 ± 1.40; CNT intermediate stage: -22.13 ± 1.10; CNT late stage: -25.08 ± 0.76; 6-OHDA early stage: -24.49 ± 0.61; 6-OHDA intermediate stage: -24.82 ± 0.45; 6-OHDA late stage: -24.82 ± 0.45. Two-way ANOVA with Sidak's multiple comparisons test $F_{StagesXGroups}$ (2, 141) = 0.50 p=0.61, $F_{Stages}$ (2, 141) = 0.46 p=0.63, $F_{Groups}$ (1, 141) = 0.45 p=0.50.

**(E)** Longitudinal cortical power of beta oscillations in controls in blue (CNT n=8) and 6-OHDA mice in orange (n=13). CNT early stage: -22.60 ± 1.46; CNT intermediate stage: -19.29 ± 0.42; CNT late stage: -20.97 ± 0.48; 6-OHDA early stage: -20.90 ± 0.49; 6-OHDA intermediate stage: -19.79 ± 0.48; 6-OHDA late stage: -20.84 ± 0.90. Two-way ANOVA with Sidak's multiple comparisons test $F_{StagesXGroups}$ (2, 163) = 1.380 p=0.25, $F_{Stages}$ (2, 163) = 6.07 p=0.0029, $F_{Groups}$= 0.45 p=0.50.

**(F)** Longitudinal cortical power of gamma oscillations in controls in blue (CNT n=8) and 6-OHDA mice in orange (n=13). CNT early stage: -30.31 ± 1.46; CNT intermediate : -29.16 ± 0.65; CNT late stage: -30.57 ± 0.73; 6-OHDA early stage: -29.85 ± 0.66; 6-OHDA intermediate stage: -30.04 ± 0.46; 6-OHDA late stage: -29.58 ± 0.56. Two-way ANOVA with Sidak's multiple comparisons test $F_{StagesXGroups}$ (2, 141) = 1.11 p=0.33, $F_{Stages}$ (2, 141) = 0.36 p=0.70, $F_{Groups}$ (1, 141) = 0.11 p=0.75.

**(G)** Heat map representing values from the correlation matrix (Pearson correlation) of analysed parameters (bottom triangle), and the significance of each pairwise comparison (top triangle: black box indicates p < 0.05) (n=5 animals for CNT group and n=6 for 6-OHDA group).

**(H-J)** Scatter plot and linear regression detailing specific correlations depicted in **G**. In particular, we show the correlation between cortical high-gamma modulation at 28DPL and

**(H)** performance in the behavioral test at 28DPL, and

**(I)** the number of $TH^+$ cells in the SNc,

**(J)** the intensity of $TH^+$ fibers in the CPu (CNT n=5, 6-OHDA n=6). r and p-values are indicated in the figures, respectively.

Data are expressed as mean ± SEM. *p<0.05; **p<0.01; ***p<0.001; ****p<0.0001.

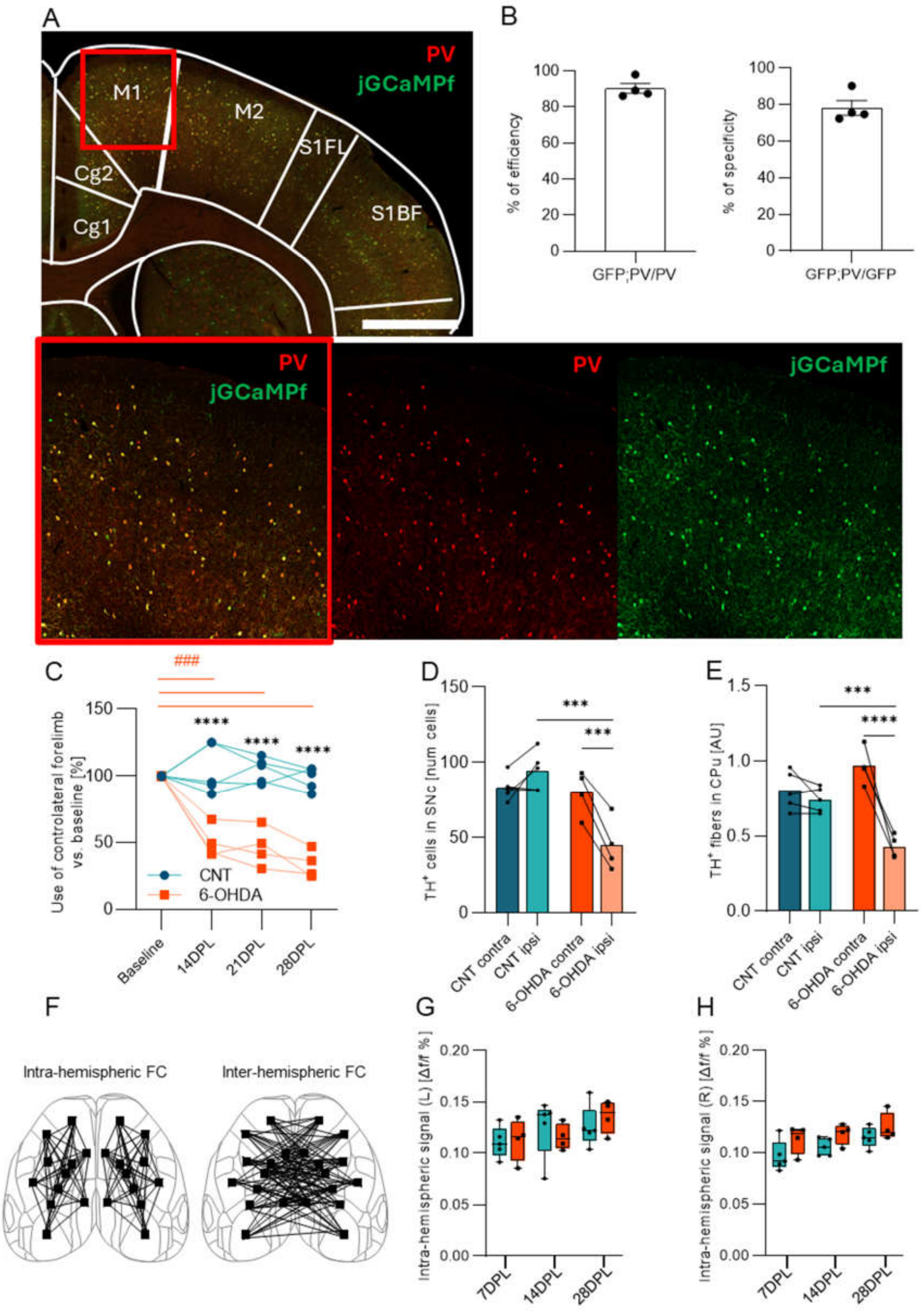


**Figure S3.**

**(A)** Representative coronal sections immunostained for PV (red) and virally expressed jGCaMPf (green). The magnification of the red box (bottom) shows M1 in merge and single channel color. Scale bar, 500 μm and 167 μm for high magnification.

**(B)** Graphs showing the efficiency (n=4, 90.2 ± 2.7 %, left) and the specificity (n=4, 78.2 ± 4.1 %, right) of viral infection in the PV-Cre mouse model.

**(C)** Forelimbs motor function was assessed using the cylinder test before 6-OHDA injection (baseline) and at 14, 21 and 28 days post lesion (14DPL, 21DPL, 28DPL). The graph illustrates the percentage of wall contacts made with the contralateral forepaw for the control group in blue (n=5, Baseline = 100.0 ± 0.0; 7DPL = 105.0 ± 3.5; 14DPL = 109.4 ± 8.0; 21DPL = 101.9 ± 4.3; 28DPL = 95.5 ± 4.1) and for 6-OHDA group in orange (n=4, Baseline = 100.0 ± 0.0; 7DPL = 51.1 ± 9.5; 14DPL = 50.6 ± 4.7; 21DPL = 62.5 ± 18.9; 28DPL = 36.3 ± 4.8). Repeated Two-way ANOVA with Sidak's multiple comparisons test $F_{TimeXGroups}$ (3, 21) = 25.14 p<0.0001, $F_{Time}$ (3, 21) = 23.39 p<0.0001, $F_{Groups}$ (1, 7) = 69.59 p<0.0001; 14DPL CNT vs 6-OHDA p<0.0001; 21DPL CNT vs 6-OHDA p<0.0001; 28DPL CNT vs 6-OHDA p<0.0001; One sample t-test with Sidak's multiple comparisons test 6-OHDA 14DPL p=0.001; 6-OHDA 21DPL p=0.001, 6-OHDA 28DPL p=0.004.

**(D)** Quantification of SNc TH$^+$ cells in both the hemisphere (contralateral and ipsilateral to the injury) for CNT in blue (n=5, contra = 83.20 ± 3.88; ipsi = 94.30 ± 5.88) and 6-OHDA in orange (n=4, contra = 80.25± 7.51; ipsi = 45.08 ± 8.65). Two-way ANOVA with Sidak's multiple comparisons test $F_{GroupsXHemisphere}$ (1, 14) = 12.96 p=0.0029, $F_{Groups}$ (1, 14) = 16.48 p=0.0012, $F_{Hemisphere}$ (1, 14) = 3.50 p=0.049; contra vs ipsi 6-OHDA p=0.0005; 6-OHDA vs CNT ipsi p=0.0002.

**(E)** Quantification of the mean fluorescence of TH$^+$ fibers relative optical density (ROD) in CPu for CNT (blue, n=5, contra = 0.80 ± 0.05; ipsi = 0.74 ± 0.03) and 6-OHDA (orange, n=4, contra = 0.96 ± 0.06; ipsi = 0.43 ± 0.04). Two-way ANOVA with Sidak's multiple comparisons test $F_{GroupsXHemisphere}$ (1, 14) = 22.17 p=0.0003, $F_{Groups}$ (1, 14) = 5.277 p=0.0338, $F_{Hemisphere}$ (1, 14) = 35.23 p<0.0001; contra vs ipsi 6-OHDA p<0.0001; 6-OHDA vs CNT ipsi p=0.001.

**(F)** Schematic representation of intra- and inter-hemispheric PV connectivity in PV-Cre mice.

**(G)** Intra-hemispheric signal of the left hemisphere, contralateral to the injury. Repeated Two-way ANOVA.

**(H)** Intra-hemispheric signal of the right hemisphere, ipsilateral to the injury. Repeated Two-way ANOVA.

In blue CNT n=6 and in orange 6-OHDA n=5. Data are expressed as mean ± SEM. *p<0.05; **p<0.01; ***p<0.001; ****p<0.0001.

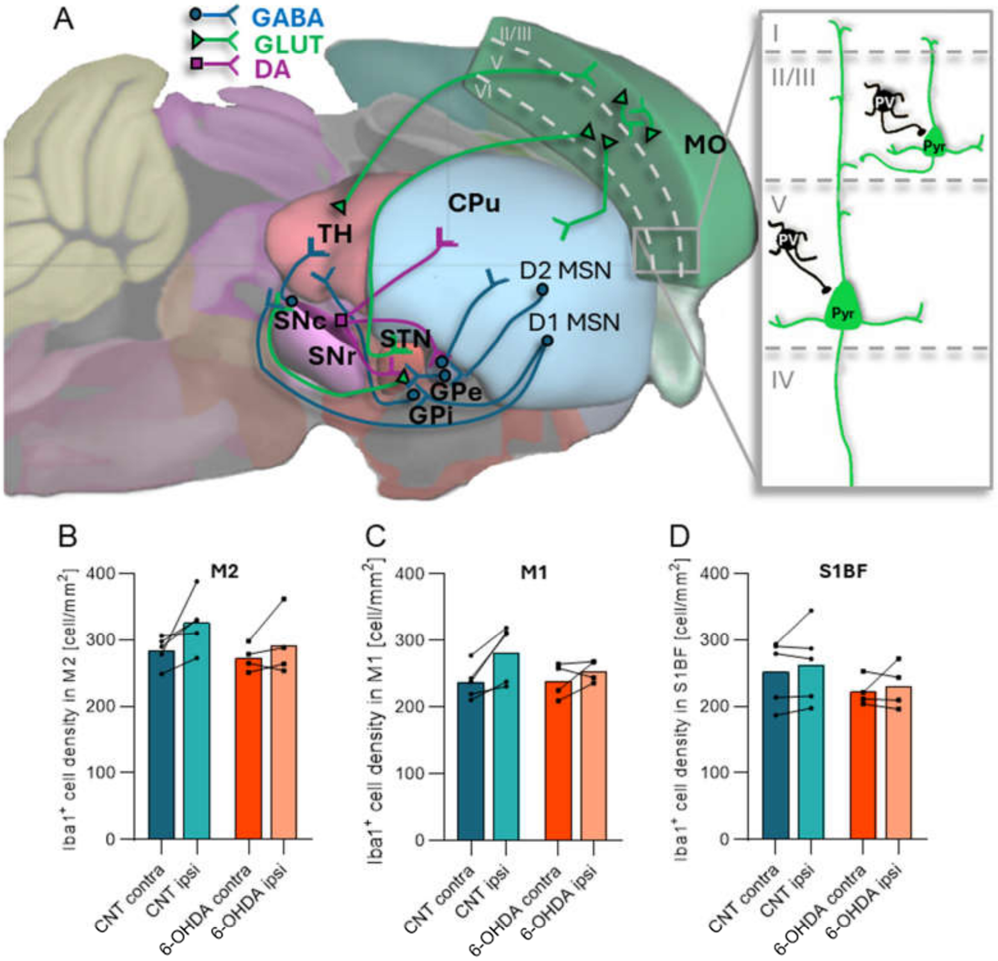

**Figure S4.**

**(A) Schematic diagram of the major pathways involved in Parkinson's Disease:** dopaminergic (in purple), glutamatergic (in green) and GABAergic (in blue) connections. **The direct pathway** facilitates movement. It is activated by dopamine from the SNc, acting on D1 receptors of GABAergic medium spiny neurons (D1 MSN) in the striatum (caudate and putamen). The striatum sends inhibitory signals to the internal globus pallidus (GPi) and the substantia nigra pars reticulata (SNr). These two regions typically inhibit the thalamus (TH), which in turn excites the somatomotor cortex (MO). **The indirect pathway** inhibits movement. In this pathway, dopamine acts on D2 receptors of GABAergic medium spiny neurons (D2 MSN) in the striatum. The striatum inhibits the external globus pallidus (GPe), which normally inhibits the subthalamic nucleus (STN). The STN excites the GPi/SNr, leading to increased inhibition of the thalamus, thus suppressing movement. In **Parkinson's disease**, the loss of dopaminergic neurons in the substantia nigra pars compacta (SNc) and their projection fibers in CPu disrupts the balance between the direct and indirect pathways in the basal ganglia, leading to motor dysfunction. **The Local Circuit of Parvalbumin-Expressing Interneurons:** PV-INs are a specialized subtype of GABAergic inhibitory neurons characterized by their fast-spiking activity. In the cortex, PV-INs play a crucial role in regulating cortical excitability and maintaining the excitatory/inhibitory (E/I) balance. By providing rapid, perisomatic inhibition to pyramidal neurons (Pyr), they precisely control the timing and synchronization of neuronal firing. This fine-tuned regulation is essential for coordinating network oscillations, particularly gamma oscillations, which are involved in cognitive processes such as perception, attention, and memory. As a result, PV-INs are key players in normal brain function and are implicated in various neurological and psychiatric disorders when dysfunctional.

**(B) PV+ cell density in secondary motor cortex cortex** (CNT contra = 284.3 ± 10.05; CNT ipsi = 326.7 ± 18.73; 6-OHDA contra = 273.3 ± 10.07; 6-OHDA ipsi = 292.1 ± 24.48), Two-way ANOVA,

**(C) in primary motor cortex** (CNT contra = 238.0 ± 11.52; CNT ipsi = 282.3 ± 19.54; 6-OHDA contra = 239.4 ± 13.20; 6-OHDA ipsi = 254.2 ± 8.180), Two-way ANOVA, and

**(D) in primary somatosensory cortex, barrel field** (CNT contra = 252.8 ± 21.96; CNT ipsi = 263.2 ± 26.36; 6-OHDA contra = 222.3 ± 10.80; 6-OHDA ipsi = 230.4 ± 17.14), Two-way ANOVA.

Data are expressed as mean ± SEM. *p<0.05; **p<0.01; ***p<0.001; ****p<0.0001

| | Early | | Intermediate | | Late | |
|---|---|---|---|---|---|---|
| | CNT | 6-OHDA | CNT | 6-OHDA | CNT | 6-OHDA |
| C) Delta power in striatum | -13.01 ± 0.55 | -12.68 ± 0.46 | -12.27 ± 0.29 | -11.72 ± 0.33 | -11.86± 0.34 | -9.76 ± 0.30 |
| E) Striatum-cortex delta coherence | 0.17 ± 0.06 | 0.14 ± 0.03 | 0.14 ± 0.03 | 0.18 ± 0.02 | 0.18 ± 0.05 | 0.30 ± 0.03 |
| F) Delta max cross corr. | 0.21 ± 0.05 | 0.19 ± 0.04 | 0.17 ± 0.03 | 0.27 ± 0.0 | 0.17 ± 0.05 | 0.37 ± 0.03 |
| G) Time lag max-cross corr. | -104.00 ± 110.24 | 17.83 ± 87.49 | -32.21 ± 83.89 | 34.60 ± 37.05 | -45.88 ± 113.10 | -11.85 ± 26.04 |
| K) Delta mod. striatum | 43.24 ± 10.83 | 25.59 ± 4.52 | 19.81 ± 6.49 | 15.89 ± 5.04 | 27.51 ± 6.50 | 7.15 ± 6.44 |
| L) High-gamma mod. striatum | 58.13 ± 10.38 | 27.35 ± 3.88 | 30.46 ± 4.48 | 18.34 ± 3.45 | 17.16 ± 3.07 | 16.46 ± 2.14 |
| M) Mean force | 0.26 ± 0.058 | 0.22 ± 0.043 | 0.24 ± 0.017 | 0.21 ± 0.022 | 0.27 ± 0.030 | 0.28 ± 0.033 |
| N) Delta mod. cortex | 73.43 ± 11.52 | 83.96 ± 11.97 | 50.35 ± 8.97 | 44.54 ± 9.76 | 24.03 ± 5.03 | 47.9 ± 9.30 |
| O) High-gamma mos. cortex | 49.12 ± 3.56 | 42.54 ± 3.16 | 35.98 ± 3.13 | 41.02 ± 3.50 | 37.91 ± 3.77 | 50.33 ± 4.04 |
| P) Area under the curve | 2.69 ± 0.38 | 2.10 ± 0.46 | 2.93 ± 0.21 | 2.84 ± 0.49 | 3.18 ± 0.30 | 5.13 ± 0.86 |

**Table S1.** Electrophysiological data values corresponding to Figure 1, grouped by experimental conditions (CNT and 6-OHDA) across different time points (early, intermediate, late).

| | 14DPL | | 21DPL | | 28DPL | |
|---|---|---|---|---|---|---|
| | CNT | 6-OHDA | CNT | 6-OHDA | CNT | 6-OHDA |
| H) Global FC | 0.042 ± 0.003 | 0.062 ± 0.006 | 0.045 ± 0.006 | 0.064 ± 0.005 | 0.043 ± 0.005 | 0.073 ± 0.003 |
| G) Inter-hem FC | −0.020 ± 0.002 | 0.012 ± 0.008 | −0.025 ± 0.008 | 0.012 ± 0.007 | −0.035 ± 0.011 | 0.016 ± 0.007 |

| | | | | | | |
|---|---|---|---|---|---|---|
| I) MOs-$p_L$ FC | −0.041 ± 0.019 | −0.046 ± 0.017 | −0.025 ± 0.009 | −0.025 ± 0.017 | −0.038 ± 0.063 | −0.021 ± 0.021 |
| J) MOs-$p_R$ FC | 0.016 ± 0.008 | 0.034 ± 0.015 | 0.021 ± 0.011 | 0.032 ± 0.005 | 0.013 ± 0.010 | 0.052 ± 0.017 |
| K) MOp-$p_L$ FC | −0.071 ± 0.012 | −0.11 ± 0.007 | −0.087 ± 0.021 | −0.087 ± 0.021 | −0.031 ± 0.034 | −0.095 ± 0.030 |
| L) MOp-$p_R$ FC | −0.081 ± 0.013 | −0.10 ± 0.032 | −0.055 ± 0.013 | −0.12 ± 0.007 | −0.047 ± 0.016 | −0.11 ± 0.010 |
| M) SSp-$BDF_L$ FC | 0.093 ± 0.013 | 0.049 ± 0.017 | 0.087 ± 0.005 0 | 0.042 ± 0.021 | 0.1 ± 0.007 | 0.027 ± 0.022 |
| N) SSp-$BDF_R$ FC | 0.064 ± 0.004 | 0.021 ± 0.015 | 0.011 ± 0.022 0 | 0.053 ± 0.022 | 0.12 ± 0.020 | 0.035 ± 0.028 |

**Table S2.** Functional connectivity (FC) values corresponding to Figure 2, grouped by experimental conditions (CNT and 6-OHDA) across different time points (early, intermediate, late).

| | CNT | | 6-OHDA | |
|---|---|---|---|---|
| | contra | ipsi | contra | ipsi |
| B) PV distribution on M2 | 374.83 ± 10.97 | 380.47 ± 17.74 | 226.78 ± 54.60 | 251.00 ± 53.29 |
| C) PV distribution on M1 | 653.72 ± 28.58 | 636.94 ± 30.20 | 354.06 ± 61.90 | 367.25 ± 52.92 |
| D) PV distribution on S1BF | 480.38 ± 33.34 | 489.38 ± 18.18 | 356.89 ± 66.34 | 414.46 ± 75.21 |
| F) PV puncta in M2 | 13.10 ± 0.81 | 13.61 ± 0.61 | 11.0 ± 0.88 | 14.30 ± 1.04 |
| G) PV puncta in M1 | 13.87 ± 0.83 | 13.53 ± 1.02 | 9.85 ± 0.34 | 16.16 ± 0.94 |
| H) PV puncta in S1BF | 16.15 ± 0.43 | 16.59 ± 0.70 | 12.14 ± 0.76 | 16.36 ± 1.44 |
| I) PV puncta/density in M2 | 0.035 ± 0.0027 | 0.036 ± 0.0020 | 0.064 ± 0.014 | 0.070 ± 0.014 |
| J) PV puncta/density in M1 | 0.034 ± 0.0043 | 0.036 ± 0.0040 | 0.042 ± 0.0.0045 | 0.076 ± 0.0096 |
| K) PV puncta/density in S1BF | 0.052 ± 0.0037 | 0.050 ± 0.0032 | 0.052 ± 0.0066 | 0.068 ± 0.0093 |
| M) PV cell intersections | 2.98 ± 0.278 | 3.03 ± 0.27 | 3.40 ± 0.25 | 3.35 ± 0.24 |
| N) PV dendrite tot branching pt | 21.12 ± 0.92 | 21.19 1.14 | 24.80 ± 1.83 | 16.46 ± 0.92 |

**Table S3.** PV-IN histological values corresponding to Figure 3, grouped by experimental conditions (CNT and 6-OHDA) across different time points (early, intermediate, late).

| Two-way ANOVA-Sidak's multiple comparisons test | M2 | | | | M1 | | | | S1BF | | | |
|---|---|---|---|---|---|---|---|---|---|---|---|---|
| | CNT contra | CNT ipsi | 6-OHDA contra | 6-OHDA ipsi | CNT contra | CNT ipsi | 6-OHDA contra | 6-OHDA ipsi | CNT contra | CNT ipsi | 6-OHDA contra | 6-OHDA ipsi |
| Bin1 vs. Bin2 | 0.0010 | <0.0001 | 0.6129 | 0.1525 | <0.0001 | <0.0001 | 0.0013 | 0.0063 | <0.0001 | <0.0001 | 0.0012 | 0.0009 |

| | | | | | | | | | | | | |
|---|---|---|---|---|---|---|---|---|---|---|---|---|
| Bin1 vs. Bin3 | 0.0002 | 0.0001 | 0.1073 | 0.0035 | <0.0001 | 0.002 | 0.0015 | 0.0071 | <0.0001 | <0.0001 | <0.0001 | 0.0009 |
| Bin1 vs. Bin4 | | | | | <0.0001 | <0.0001 | 0.0010 | 0.0005 | <0.0001 | <0.0001 | 0.0004 | 0.0008 |
| Bin1 vs. Bin5 | | | | | 0.7386 | 0.9696 | 0.9821 | >0.9999 | 0.9704 | 0.6907 | >0.9999 | 0.9945 |
| Bin2 vs. Bin3 | 0.8612 | 0.9755 | 0.9755 | 0.2730 | 0.9993 | 0.4419 | >0.9999 | >0,9999 | 0.9525 | 0.9983 | 0.9399 | >0.9999 |
| Bin2 vs. Bin4 | | | | | 0.9952 | 0.8878 | >0.9999 | 0.9424 | 0.9855 | 0.9983 | 0.9984 | >0.9999 |
| Bin2 vs. Bin5 | | | | | <0.0001 | <0.0001 | 0.0074 | 0.0083 | <0.0001 | <0.0001 | 0.0010 | 0.0002 |
| Bin3 vs. Bin4 | | | | | 0.9716 | 0.0778 | >0.9999 | 0.9322 | 0.9996 | 0.9437 | 0.9894 | >0.9999 |
| Bin3 vs. Bin5 | | | | | <0.0001 | <0.0001 | 0.0086 | 0.0092 | <0.0001 | <0.0001 | <0.0001 | 0.0002 |
| Bin4 vs. Bin5 | | | | | <0.0001 | <0.0001 | 0.0057 | 0.0007 | <0.0001 | <0.0001 | 0.0004 | 0.0002 |

| Two-way ANOVA-Sidak's multiple comparisons test | M2 | | | M1 | | | | | S1BF | | | | |
|---|---|---|---|---|---|---|---|---|---|---|---|---|---|
| | Bin 1 | Bin 2 | Bin 3 | Bin 1 | Bin 2 | Bin 3 | Bin 4 | Bin 5 | Bin 1 | Bin 2 | Bin 3 | Bin 4 | Bin 5 |
| CNT contra vs. CNT ipsi | >0.9999 | 0.7977 | 0.9979 | 0.9995 | 0.7986 | 0.9816 | 0.9816 | 0.8953 | 0.9760 | 0.8795 | 0.9629 | 0.9999 | >0.9999 |
| CNT contra vs. 6-OHDA contra | 0.0724 | >0.0001 | 0.0003 | 0.0292 | >0.0001 | >0.0001 | >0.0001 | 0.6815 | 0.4853 | 0.1297 | 0.1693 | 0.0877 | >0.9999 |
| CNT ipsi vs. 6-OHDA ipsi | 0.0695 | >0.0001 | 0.0129 | 0.0528 | >0.0001 | >0.0001 | >0.0001 | 0.2283 | 0.8489 | 0.2713 | 0.1204 | 0.4451 | >0.9999 |
| 6-OHDA contra vs. 6-OHDA ipsi | >0.9999 | 0.7839 | 0.5215 | 0.9884 | 0.9982 | 0.9590 | 0.9590 | 0.9991 | 0.7743 | 0.7299 | 0.9929 | 0.8592 | >0.9999 |

**Table S4.** Statistics of PV density and distribution data corresponding to Figure 3 B, C, D. Highlighted in yellow are significant values p<0.05.

| | CNT | | 6-OHDA | |
|---|---|---|---|---|
| | contra | ipsi | contra | ipsi |
| B) VGAT puncta in M1 | 10.51 ± 0.38 | 11.29 ± 0.93 | 9.02 ± 0.29 | 9.82 ± 0.23 |
| D) VGLUT1 puncta in M1 | 22.55 ± 0.56 | 23.24 ± 0.35 | 20.95 ± 0.56 | 27.49 ± 0.56 |
| F) VGLUT2 puncta in M1 | 4.32 ± 0.34 | 4.54 ± 0.45 | 5.10 ± 0.17 | 8.22 ± 0.20 |

| | | | | |
|---|---|---|---|---|
| H) CD68$^+$ in Iba1$^+$ | 9.33 ± 0.84 | 12.02 ± 1.07 | 12.00 ± 0.83 | 22.41 ± 2.00 |
| I) VGLUT1 in CD68$^+$/Iba1$^+$ | 0.70 ± 0.086 | 0.74 ± 0.062 | 0.79 ± 0.063 | 0.96 ± 0.062 |

**Table S5.** Histological data values corresponding to Figure 5, grouped by experimental conditions (CNT and 6-OHDA) across different time points (early, intermediate, late).

**Abbreviations**

| | |
|---|---|
| 6-OHDA | 6-hydroxydopamine |
| E/I | excitatory/inhibitory balance |
| FC | functional connectivity |
| NGF | nerve growth factor |
| PD | Parkinson's disease |
| PV-IN | parvalbumin-positive interneuron |
| tACS | transcranial alternating current stimulation |
| TH | tyrosine hydroxylase |
| VGAT | vesicular GABA transporter |
| VGLUT1 | vesicular glutamate transporter type 1 |
| VGLUT2 | vesicular glutamate transporter type 2 |
| WF | Wide-Field |

Brain regions:

| | |
|---|---|
| CPu | striatum |
| M1 | primary motor cortex |
| M2 | secondary motor cortex |
| MO | motor cortex |
| GPe | external globus pallidus |
| GPi | internal globus pallidus |
| S1BF | primary somatosensory cortex barrel field |
| SNc | substantia nigra pars compacta |
| SNr | substantia nigra pars reticulata |
| STN | subthalamic nucleus |
| TH | thalamus |

WF ROIs:

| | |
|---|---|
| MOs-a$_L$ | secondary motor area, anterior domain (left hemisphere) |
| MOs-p$_L$ | secondary motor area, posterior domain (left hemisphere) |
| MOp-a$_L$ | primary motor area, anterior domain (left hemisphere) |
| MOp-p$_L$ | primary motor area, posterior domain (left hemisphere) |
| SSp-bfd$_L$ | primary somatosensory area (left hemisphere) |
| SSp-tr$_L$ | primary somatosensory area, trunk region (left hemisphere) |
| SSp-fl$_L$ | primary somatosensory area, forelimb region (left hemisphere) |
| SSp-hl$_L$ | primary somatosensory area, hindlimb region (left hemisphere) |
| RSP$_L$ | retrosplenial area (left hemisphere) |
| VISa$_L$ | anterior visual area (left hemisphere) |
| VISp$_L$ | posterior visual area (left hemisphere) |
| MOs-a$_R$ | secondary motor area, anterior domain(right hemisphere) |
| MOs-p$_R$ | secondary motor area, posterior domain (right hemisphere) |
| MOp-a$_R$ | primary motor area, anterior domain (right hemisphere) |
| MOp-p$_R$ | primary motor area, posterior domain (right hemisphere) |
| SSp-bfd$_R$ | primary somatosensory area, barrel field (right hemisphere) |
| SSp-tr$_R$ | primary somatosensory area, trunk domain (right hemisphere) |
| SSp-fl$_R$ | primary somatosensory area, forelimb region (right hemisphere) |
| SSp-hl$_R$ | primary somatosensory area, hindlimb region (right hemisphere) |
| RSP$_R$ | retrosplenial area (right hemisphere) |
| VISa$_R$ | anterior visual area (right hemisphere) |

| | |
|---|---|
| $VISp_R$ | posterior visual area |
